# Multiscale effects of perturbed translation dynamics inform antimalarial design

**DOI:** 10.1101/2023.09.03.556115

**Authors:** Leonie Anton, Wenjing Cheng, Meseret T. Haile, David W. Cobb, Xiyan Zhu, Leyan Han, Emerson Li, Anjali Nair, Carolyn L. Lee, Hangjun Ke, Guoan Zhang, Emma H. Doud, Chi-Min Ho

## Abstract

Malaria parasites rely heavily on rapid, high fidelity protein synthesis to infect and replicate in human erythrocytes, making translation an attractive target for new antimalarials. Here, we have determined *in situ* structures of *Pf*80S ribosomes in thirteen conformational and compositional states from cryoFIB-milled *Plasmodium falciparum*-infected human erythrocytes across the stages of asexual intraerythrocytic parasite replication. We observe eight active translation intermediates, enabling us to define the native malarial translation elongation cycle, which surprisingly features a bifurcation at the decoding stage of the cycle that has not previously been described. Examination of perturbations in the distribution of ribosomes among these states in the presence of a malaria-specific translation inhibitor suggests that the inhibitor impedes *Pf*eEF2 and *Pf*eEF1α interactions with the ribosome. We integrated our *in situ* cryoET data with proteomic and ultrastructural data to arrive at a deeper understanding of malarial translation, which will inform development of new therapies.

## INTRODUCTION

Malaria exacts a heavy toll on global health, with half the world’s population currently at risk^1^. In the face of setbacks during the covid-19 pandemic^1^ and the continued rise of artemisinin-resistance^2^, new treatments with novel modes of action are urgently needed. *Plasmodium* parasites, the causative agents of malaria, invade and replicate inside human red blood cells, directly leading to morbidity and mortality^3,4^. Upon entering host erythrocytes, malaria parasites adhere to a strict schedule of growth and asexual replication, progressing through four morphologically distinct stages every 48-hours (Figure 1A, B). Each cycle starts with newly-formed ring-stage parasites directly after invasion, then progresses through the trophozoite and schizont stages, culminating in 16-32 daughter cells called merozoites, which lyse the host cell, egress, and reinvade new uninfected erythrocytes to restart the cycle^3,4^. Progression of the parasites through these morphologically and functionally distinct stages is precisely timed, requiring tightly coordinated activation and deactivation of genes to achieve the strictly prescribed bloc of gene expression profiles required for each life cycle stage^5–7^. To accomplish this, rapid, large-scale mRNA transcription and protein synthesis is carried out in a tightly choreographed “just-in-time” model where genes are transcribed and proteins made immediately before they are needed^5–8^. As the key executor of this process, the translational machinery of the malaria parasite has emerged as an attractive target for therapeutic intervention and is directly implicated in the mode of action of a leading antimalarial therapy currently in clinical trials^9^.

**Figure 1.**
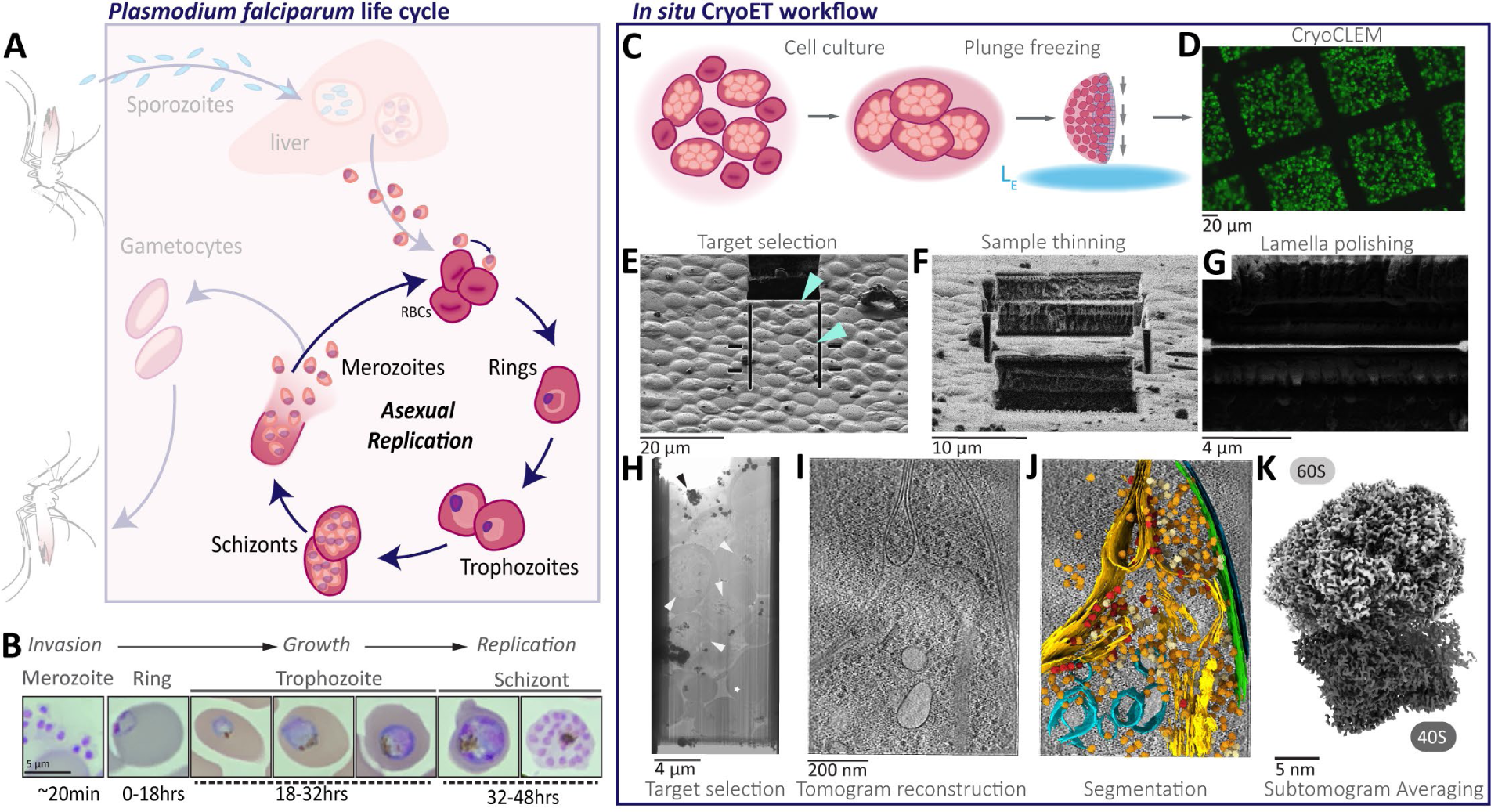
Visualizing molecular details of *P. falciparum* asexual intraerythrocytic life cycle stages using *in situ* cryo electron tomography (cryoET). **A**, Schematic of the of *P. falciparum* life cycle, with focus on the intraerythrocytic asexual replication stages in human red blood cells (RBCs). **B**, Giemsa-stained, parasite-infected RBCs (iRBCs) at the different intraerythrocytic stages from invasion to replication. **C-J,** *In situ* cryoET workflow from sample preparation to data analysis. **C**, iRBCs cultured in suspension are applied to a grid support and plunge frozen in liquid ethane (L_E_). **D**, Cryo correlative light and electron microscopy (cryoCLEM) image of frozen grids showing distribution of fluorescently labelled parasites. **E**, Top down view of a frozen grid in a cryo focused-ion beam scanning electron microscope (cryoFIB-SEM). Arrow heads indicate schizont infected RBCs. **F-G**, FIB images of the sample thinned to ∼3µm (**F**) and the final thinned cross-section (lamella) at ∼150nm (**G**), taken at the milling angle. **H**, Montaged cryo transmission electron microscope (cryoTEM) image of the entire lamella. Parasites are indicated with white arrow heads, contamination with black arrow heads and milling artifacts with a white star. **I**, Averaged central slice of a tomographic volume reconstructed from tilt series collected on lamella. **J**, Ultrastructural segmentations and subtomogram averaged *Pf*80S ribosomes mapped back into the tomographic volume and overlaid onto the corresponding central slice from (**I**). **K**, Final 4.1Å resolution subtomogram averaged consensus reconstruction of *Pf*80S.

Unfortunately, our understanding of the translational machinery and its regulation in *Plasmodium* is limited compared to our knowledge of protein synthesis in bacteria, yeast, or humans, hindering targeted development of new *Plasmodium*-specific translation inhibitors. This is due in part to a paucity of structural information, as the *P. falciparum* proteome has long resisted structural study. As such, our knowledge is currently limited to a handful of published single particle cryo electron microscopy (cryoEM) structures from purified *Pf*80S ribosomes^10,11^, which do not represent the full ensemble of ribosomal states in the cell, nor elucidate the distribution of ribosomes among these states.

In this study, we have resolved *in situ* structures of the *Pf*80S ribosome from cryo-preserved *P. falciparum*-infected human erythrocytes across the asexual intraerythroctyic stages. We then used a multi-disciplinary approach to define how inhibiting translation perturbs the distribution of ribosomal states in the parasite and establish the consequent effects on the molecular, ultrastructural and cellular integrity of the parasite. This is the first *in situ* study of ribosomes from an organism that is evolutionarily very distant from commonly studied model organisms and provides significant new insights into the broad range of ribosome function throughout the eukaryotic lineage.

## RESULTS

### Consensus structure of the *Pf*80S ribosome in the native cellular milieu

To determine native structures of the *P. falciparum* translation machinery using *in situ* cryo electron tomography (cryoET), we used a cryo-focused ion beam scanning electron microscope (cryoFIB-SEM) to create 150 - 200nm thin cellular sections (lamellae) of cryo-preserved *P. falciparum* parasites isolated at the trophozoite, schizont and merozoite stages from synchronous *in vitro* cultures (Figure 1C-F). As ring-stage parasites are dwarfed by the enveloping host erythrocyte and occupy a very small percentage of the total host-cell volume, we were unable to obtain sufficient *in situ* data for downstream analysis from this stage. To get as close to the native state as possible, we prepared frozen grids containing ribosomes crudely enriched from ring-stage parasites cultured *in vitro*, eschewing established ribosome purification schemes to minimize potential loss of native conformational states and binding partners during purification. We amassed large datasets of tomograms reconstructed from cryoET tilt series collected on the lamella (*in situ* trophozoites, schizonts, merozoites) or frozen lysates (*ex vivo* ring-stage parasites). In total, we collected 403 tomograms from 40 lamella containing 173 cells in addition to 480 tomograms from the *ex vivo* ring-stage sample (Table S1). We then identified and extracted *Pf*80S ribosome particles from all the tomograms and performed subtomogram averaging and three-dimensional (3D) refinement on the combined particles, yielding a consensus structure at an overall resolution of 4.1 Å (Figure 2A, B). Resolutions reach the Nyquist limit of 3.4 Å in the core of the large subunit and range from 3.4 - 4.4 Å throughout most of the map (Figure 2B, S1A), with some lower resolution (5 - 10 Å) areas at the periphery, particularly in the head region of the 40S small subunit, where the inherent flexibility of this part of the ribosome often leads to poorer resolutions in consensus maps (Figure 2B, S1B). The overall conformational and compositional state of the consensus map is consistent with previously published *in vitro* structures of the *Pf*80S ribosome determined by single-particle cryoEM^10,11^.

**Figure 2.**
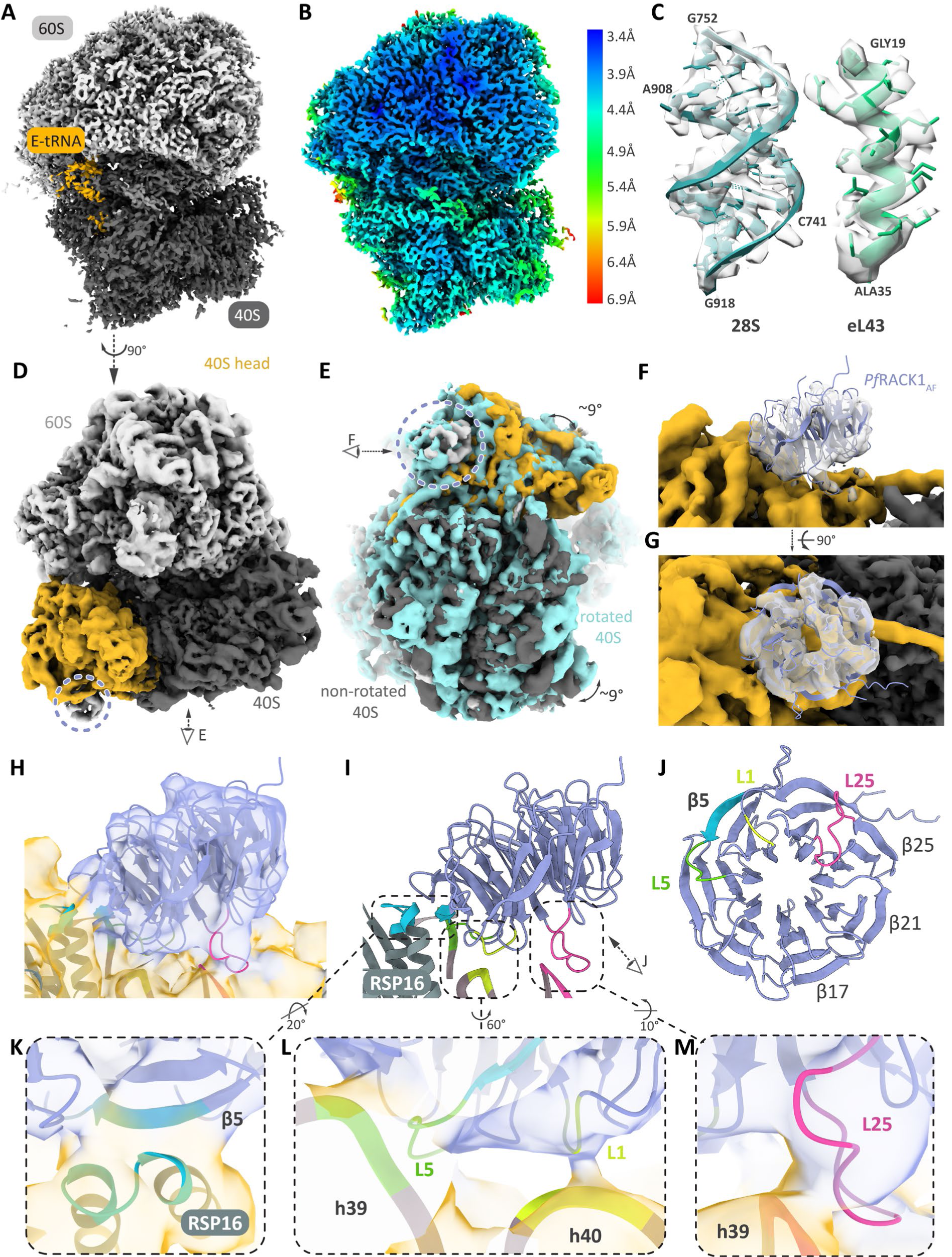
High resolution details of the native *Pf*80S ribosome determined by *in situ* cryoET. **A,** Subtomogram averaged (STA) consensus reconstruction of the *Pf*80Sribosome at an overall resolution of 4.1Å. **B,** *Pf*80S consensus map, shown in cross-section and colored according to local resolution, as calculated in RELION3.1. Majority of the map exhibits resolutions between 3.4 (Nyquist) - 4.4 Å. **C,** Ribbon models of 28S rRNA and eL43 ribosomal protein segments, shown with corresponding STA density (surface representation). **D,** Non-rotated *Pf*80S (nrt80S) consensus reconstruction. Lavender circle indicates *Pf*RACK1 in 40S head region (light orange). **E,** Overlaid top view of rotated 80S (rt80S, cyan) and nrt80S. View indicated by eye icon in **(D)**. **F-G,** Side **(F)** and top **(G)** views of AlphaFold2 *Pf*RACK1_AF_ model fit into corresponding density on 40S head region. View in **(F)** indicated by eye icon in **(E)**. **H,** Magnified view of *_Pf_*RACK1_AF_ model fit in 40S head region. Map is shown at 50% transparency and models for ribosomal protein RSP16 and relevant rRNA are shown as ribbons. **I,** M_od_el of *Pf*RACK1_AF_ binding to RSP16 and rRNA helices 39 (h39) and 40 (h40). Loop 25 knob (L25) and previously proposed interaction regions are indicated with dotted lines and colored as follows: L25 knob in deep pink, RSP16-β5 in light blue, loops 1 (L1) and 5 (L5) binding to h40 (yellow) and h39 (green), respectively. **J,** *Pf*RACK1_AF_, viewed from the 40S binding interface, with the L25 and proposed binding regions color coded. **K-M**, Detailed views of regions indicated in (**I**). Rotations from view in **(I)** are indicated.

### *Pf*RACK1 is bound to the *Pf*80S ribosome across all life cycle stages

Surprisingly, we observe a strong density in the 40S head region, which was missing in all previously published single-particle cryoEM structures of the *Pf*80S ribosome^10,11^. This location is strongly conserved as the binding site for the receptor for activated C kinase 1 (RACK1), a well-known molecular scaffold upon which translational modulators dock to modulate the activity of the ribosome^12^. RACK1 has been observed attached to the head region of the 40S small subunit in this location in other eukaryotic ribosome structures^13,14,15,16.^

As such, the conspicuous absence of *Pf*RACK1 in all high resolution single-particle cryoEM studies of the *Pf*80S ribosome to date was the source of much excitement in the field, as major points of divergence between the *Plasmodium* and human translation machineries represent areas of opportunity for therapeutic intervention. Initial speculation that RACK1 could play a very different role in *Plasmodium^11^*, ^17^ was disproven by subsequent genetic knock-down studies demonstrating that *Pf*RACK1 not only plays a key role in translational regulation, but is in fact essential for parasite growth and survival in host erythrocytes^18^. This led to further speculation that *Pf*RACK1 might function in translational regulation without interacting with the ribosome, or that *Pf*RACK1 interaction with the ribosome could vary depending on the life cycle stage or monosome versus polysome status^18^.

These speculations can now be put to rest, as we observe *Pf*RACK1 uniformly associated with the head of the *P. falciparum* 40S ribosomal small subunit across all life-cycle stages (Figure S1D). Focused classification to separate rotated and non-rotated conformations of the flexible 40S small subunit yielded a higher resolution map of the 40S at 4.3 Å overall (Figure 2D, E), with *Pf*RACK1 density resolved to sufficient quality to enable rigid body fitting of a model of *Pf*RACK1 generated using AlphaFold^19^ (*Pf*RACK1_AF_, AF-Q8IBA0-F1) (Figure 2F-J). In our map, density is visible corresponding to β-sheet 5 and the 1 and 5 loops, which form the basis of RACK1 interaction with the ribosomal protein bS16 (RSP16) and 18S helix 40 and 39 of the 40S subunit in homologs, respectively (Figure 2I-L). Density for the canonical RACK1 “knob” loop 25 is also visible in the expected location, in close proximity with h39 [^20^] (Figure 2I, M).

### Native malarial translation elongation cycle incorporates a unique combination of intermediates

The native translational landscape is defined by the full ensemble of compositional and conformational states sampled by cellular ribosomes and the distribution of ribosomes among these states within the native cellular context. To obtain sufficient numbers of *Pf*80S particles to allow extensive classification into the many ribosomal states in the parasite, we collected additional datasets and then performed multiple rounds of 3D classification to separate out the different ribosomal states (Table 1). Focused classification with masks around the tRNA and GTPase binding sites of the ribosome yielded a total of 13 different ribosomal states, comprising ∼57% of our dataset (Figure 3, Figure S1H, S2A, S4A). Around 18% of ribosomes could not be identified as a specific state (Figure S2C, S4A) and an additional 25% of our datasets sorted into lower resolution classes that could not be readily assigned to any known ribosomal states (Figure S2D, S4A). Comparison with previously published ribosomal structures from *P. falciparum* and other species enabled us to assign 8 of our high resolution states to distinct steps in translation elongation, revealing 8 active translation intermediates of the *P. falciparum* translation elongation cycle^10,11,21–24^ (Figure 3A, S1E, S2, S3A, C, E-J, S4A-C), as well as three hibernating *Pf*80S states, in the native cellular context (Figure 3B, S4D-F). The eEF2 hibernating-1 and -2 states (eEF2, E and eEF2 only) have been previously described^24,25^(Figure 3B, S1F, S4E-F). The eEF1α, E state has not previously been described, but based on its similarity to the eEF2 hibernating-1 state, we suspect this is also a hibernating state (Figure 3B, S4D). The twelfth and least populated state contains an eEF2 in the GTPase site and a density in the P site that is likely a tRNA (Figure S1G, S4B). The thirteenth state, which we have termed the unloaded state, contains nothing but an E-site tRNA and does not fit into the classical translational cycle (Figure 3B, Figure S3J). We see strong nascent peptide chain density in several of our translation intermediates (Figure S1E), but the density in this area is much weaker in the unloaded state, suggesting it is inactive (Figure S1E). The unloaded state has previously been observed in single-particle cryoEM studies^10,11^ and while its precise function is still unclear, its prevalence in our *in situ* samples suggests some physiological relevance. It is possible that this represents an idle or storage ribosome state, although we do not observe density corresponding with any known ribosome-inactivating factors in our map^26^. Furthermore the single tRNA is bound in the canonical E-site, excluding the possibility of a Z-site tRNA-associated inactive state^27,25^.

**Figure 3.**
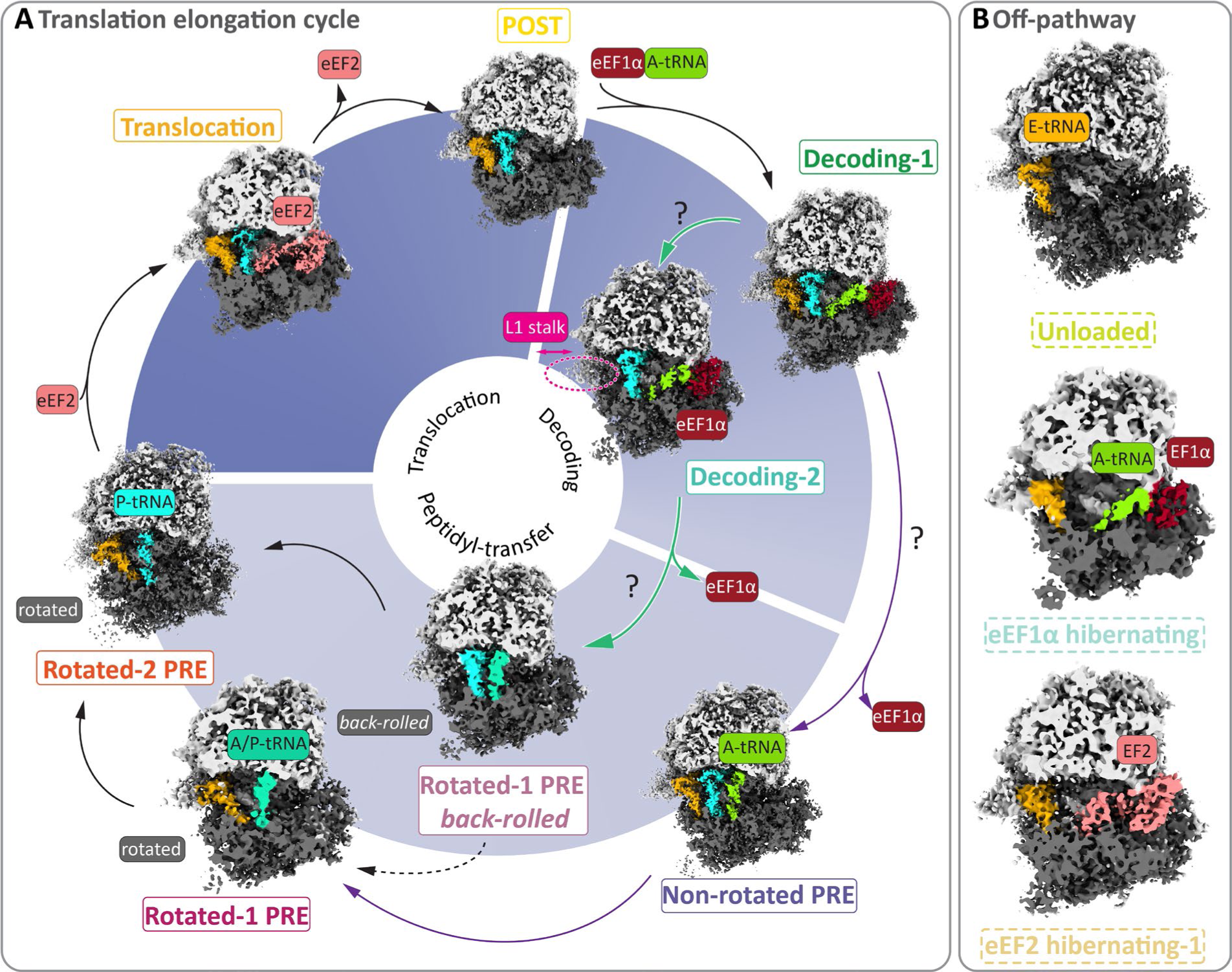
Ensemble of *in situ Pf*80S ribosomes comprised of translation intermediates and off-pathway states. **A-B,** Native structures of the *P. falciparum* translational machinery, determined by *in situ* cryoET, elucidate the molecular ensemble of *Pf*80S ribosomes in malaria parasites. **A-B,** Translation intermediate states **(A)** and off-pathway ribosomal states **(B)** by position of principal ligands in these high resolution subtomogram averaged reconstructions enabled reconstitution of the *P. falciparum* translation elongation cycle. Refined map for each state is reconstructed from all particles corresponding to a particular class across datasets (Figure S2A, Figure S4A, Table S2). Maps shown in cross-section for clarity. eEF1α elongation factor (dark red), A-tRNA (green), L1 stalk movement (pink), A/P-tRNA (teal) P-tRNA (cyan), eEF2 elongation factor (salmon), E-tRNA (orange).

**Table 1:**
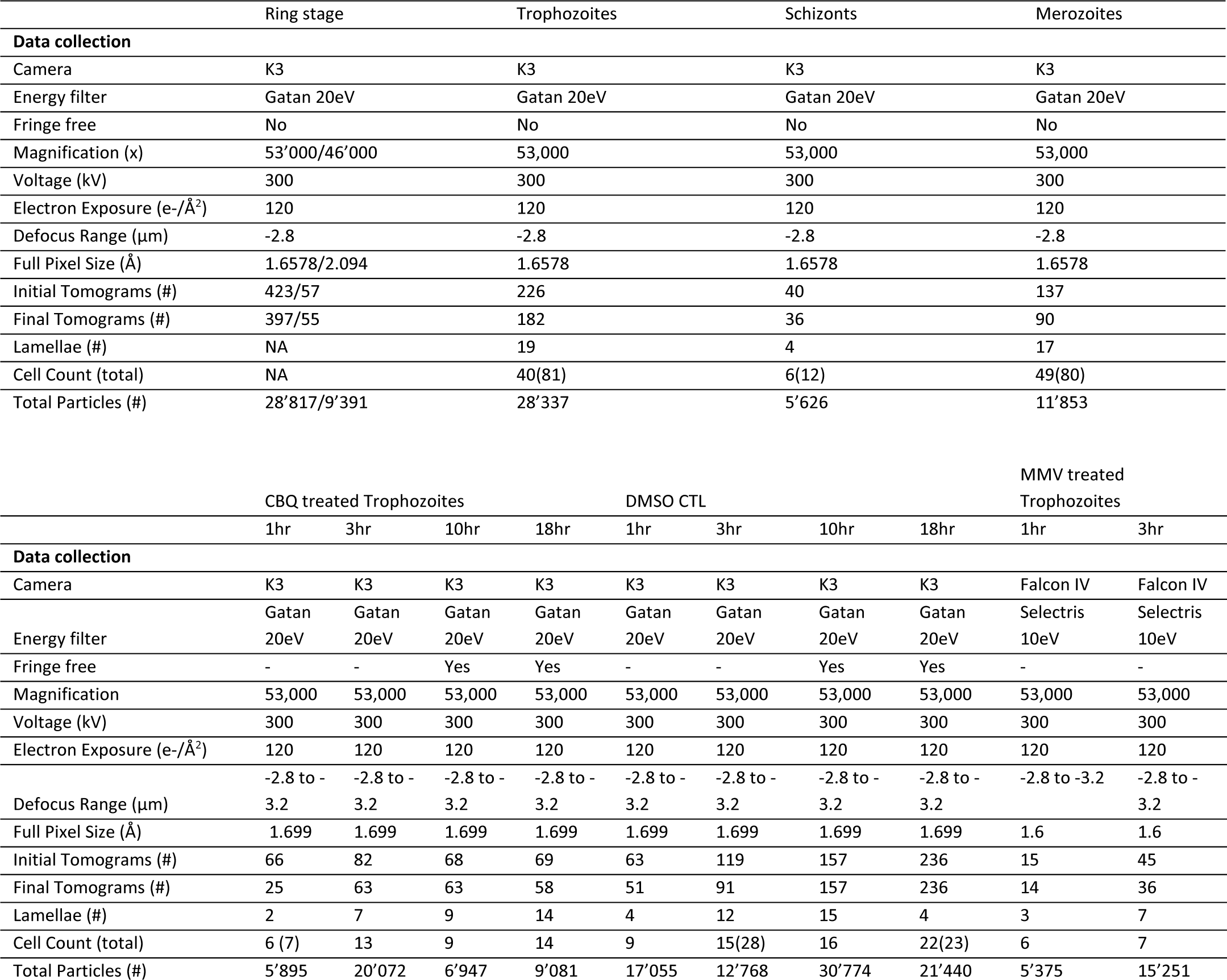
Data Collection.

|  | Ring stage | Trophozoites | Schizonts | Merozoites |
| --- | --- | --- | --- | --- |
| **Data collection** |  |  |  |  |
| Camera | K3 | K3 | K3 | K3 |
| Energy filter | Gatan 20eV | Gatan 20eV | Gatan 20eV | Gatan 20eV |
| Fringe free | No | No | No | No |
| Magnification (x) | 53'000/46'000 | 53,000 | 53,000 | 53,000 |
| Voltage (kV) | 300 | 300 | 300 | 300 |
| Electron Exposure (e-/Å<sup>2</sup>) | 120 | 120 | 120 | 120 |
| Defocus Range (µm) | -2.8 | -2.8 | -2.8 | -2.8 |
| Full Pixel Size (Å) | 1.6578/2.094 | 1.6578 | 1.6578 | 1.6578 |
| Initial Tomograms (#) | 423/57 | 226 | 40 | 137 |
| Final Tomograms (#) | 397/55 | 182 | 36 | 90 |
| Lamellae (#) | NA | 19 | 4 | 17 |
| Cell Count (total) | NA | 40(81) | 6(12) | 49(80) |
| Total Particles (#) | 28'817/9'391 | 28'337 | 5'626 | 11'853 |

|  | CBQ treated Trophozoites |  |  |  | DMSO CTL |  |  |  | MMV treated Trophozoites |  |
| --- | --- | --- | --- | --- | --- | --- | --- | --- | --- | --- |
|  | 1hr | 3hr | 10hr | 18hr | 1hr | 3hr | 10hr | 18hr | 1hr | 3hr |
| **Data collection** |  |  |  |  |  |  |  |  |  |  |
| Camera | K3 | K3 | K3 | K3 | K3 | K3 | K3 | K3 | Falcon IV | Falcon IV |
| Energy filter | Gatan 20eV | Gatan 20eV | Gatan 20eV | Gatan 20eV | Gatan 20eV | Gatan 20eV | Gatan 20eV | Gatan 20eV | Selectris 10eV | Selectris 10eV |
| Fringe free | - | - | Yes | Yes | - | - | Yes | Yes | - | - |
| Magnification | 53,000 | 53,000 | 53,000 | 53,000 | 53,000 | 53,000 | 53,000 | 53,000 | 53,000 | 53,000 |
| Voltage (kV) | 300 | 300 | 300 | 300 | 300 | 300 | 300 | 300 | 300 | 300 |
| Electron Exposure (e-/Å<sup>2</sup>) | 120 | 120 | 120 | 120 | 120 | 120 | 120 | 120 | 120 | 120 |
| Defocus Range (µm) | -2.8 to -3.2 | -2.8 to -3.2 | -2.8 to -3.2 | -2.8 to -3.2 | -2.8 to -3.2 | -2.8 to -3.2 | -2.8 to -3.2 | -2.8 to -3.2 | -2.8 to -3.2 | -2.8 to -3.2 |
| Full Pixel Size (Å) | 1.699 | 1.699 | 1.699 | 1.699 | 1.699 | 1.699 | 1.699 | 1.699 | 1.6 | 1.6 |
| Initial Tomograms (#) | 66 | 82 | 68 | 69 | 63 | 119 | 157 | 236 | 15 | 45 |
| Final Tomograms (#) | 25 | 63 | 63 | 58 | 51 | 91 | 157 | 236 | 14 | 36 |
| Lamellae (#) | 2 | 7 | 9 | 14 | 4 | 12 | 15 | 4 | 3 | 7 |
| Cell Count (total) | 6 (7) | 13 | 9 | 14 | 9 | 15(28) | 16 | 22(23) | 6 | 7 |
| Total Particles (#) | 5'895 | 20'072 | 6'947 | 9'081 | 17'055 | 12'768 | 30'774 | 21'440 | 5'375 | 15'251 |

As illustrated in Figure 3A, our datasets contain classes from all three phases corresponding with the rotated-2 PRE (P,E)^21^, translocation (eEF2, P*, E)^24,28^, POST (P*, E)^28^, decoding-1 (eEF1α, A/T, P*, E)^28^, decoding-2 (eEF1α, A/T, P*)^15^, non-rotated PRE (A, P*, E)^28^, rotated-1 PRE (A/P, E)^21^ and rotated-1 PRE *back-rolled* (A/P, P*)^25^ states of the translation elongation cycle^29^. Only the POST and rotated-2 PRE states have been characterized before in *P. falciparum* single-particle cryoEM studies^10,11^. Our decoding-1 and -2 states both contain an eEF1α-A/T-tRNA and P-site tRNA bound in a similar fashion to published decoding structures of rabbit eEF1α-A-bound 80S ribosomes, with the decoding-1 containing an E-site tRNA as well^21^ (Figure 3A, Figure S3A-D). Our decoding-2 state is consistent with the eEF1α, A/T, P class found in the protozoan *D. discoideum^22^* (Figure 3A, Figure S3C), and an AlphaFold model of *Pf*eEF1α (*Pf*eEF1α _AF_, AF-QI80P6) fits well into the density at the GTPase binding site in both decoding state reconstructions (Figure S3B,D). Our non-rotated PRE (corresponding to the classical iPRE state described in previous studies), rotated-1 PRE, rotated-1 PRE *back-rolled*, rotated-2 PRE, translocation and POST states (Figure 3A) all match previously published human and *P. falciparum* 80S single-particle cryoEM structures^11,21,23,24,25^ (Figure S3E-I, S4C). Our translocation state map contains a *Pf*eEF2 bound in an extended conformation as seen in published human or yeast ribosome-bound eEF2 structures (Figure S3H, K)^24^; ^30^. To generate a model for *Pf*eEF2 with the extended C-terminal domain, we used SWISS-MODEL with the human eEF2-bound 80S structure (PDBID:6z6m) as a template^31–35^ (Figure S3L).

Notably, comparison with recent *in situ* studies of other organisms reveals that the molecular ensemble of *Pf*80S states *in situ* comprises a novel combination of translation intermediates that has not previously been described. As of this study, translation elongation cycles follow either a decoding-1, decoding-2, rotated-1 PRE *back-rolled* and rotated-2 PRE sequence^15,22,25^ or a decoding-1, non-rotated PRE, rotated-1 PRE and rotated-2 PRE sequence for peptidyl-transfer^21,36^. The decoding-2 and rotated-1 PRE *back-rolled* states have never been seen with the non-rotated PRE state in the same cell-type before.

### *P. falciparum*-specific translation inhibitors stall parasite growth and perturb the distribution of *Pf*80S particles across ribosomal states in the cell

To explore how malaria-specific translation inhibitors might perturb the translational landscape, we analyzed our *in situ* datasets of parasites treated with *P. falciparum*-specific translation inhibitors across multiple timepoints and determined the ensemble of structures of the *Pf*80S ribosome in the native cellular context for each timepoint (Table S1). The first, known as cabamiquine^37,38,39^ (CBQ), is a leading compound currently in phase II clinical trials that has been shown to target the translation elongation factor *Pf*eEF2. The second, MMV019189, was identified as the top hit in a high-throughput screen for malarial translation inhibitors from the Medicines for Malaria Venture Pandemic and Pathogen Box^40^.

For these studies, we focused on trophozoite stage parasites, as they are the most translationally active^5,7,41^. Baragaña et al demonstrated that parasites die rapidly after exposure to CBQ beyond 48 hours, but the effects of CBQ are reversible within the first 24-48 hours of exposure if the drug is washed out^37^. To identify key time points within the first 48 hours of drug exposure for *in situ* cryoET imaging, we collected a 48-hour time course for each drug on early trophozoite-stage parasites enriched from tightly synchronized *in vitro* cultures. We incubated parasite-infected erythrocytes with drug at 20x EC50 and collected Giemsa-stained blood smears at 1, 3, and every 4^th^ hour over the 48-hour time course (Figure 4A). Control parasites successfully progressed into schizonts by 24 hours post-treatment, and had divided, egressed and re-invaded into new erythrocytes by 32 hours post-treatment. Conversely, the drug treated parasites appeared to stall at around 3 hours post-treatment, staying in that state until pyknotic cells started to appear at 20 hours post-treatment (Figure 4A,B). The percentage of pyknotic cells increased over the next 28 hours, reaching 68.6 – 76.5% by 48 hours post-treatment. Meanwhile, MMV019189 elicited a weaker growth defect than CBQ, as evidenced by a lower percentage of pyknotic cells at later time points (Figure S4G). Cycloheximide (CHX), a widely used translation inhibitor known to disrupt eEF2-mediated translocation during translation elongation^42–44^, was included as a positive control and elicited a similar phenotype as CBQ.

**Figure 4.**
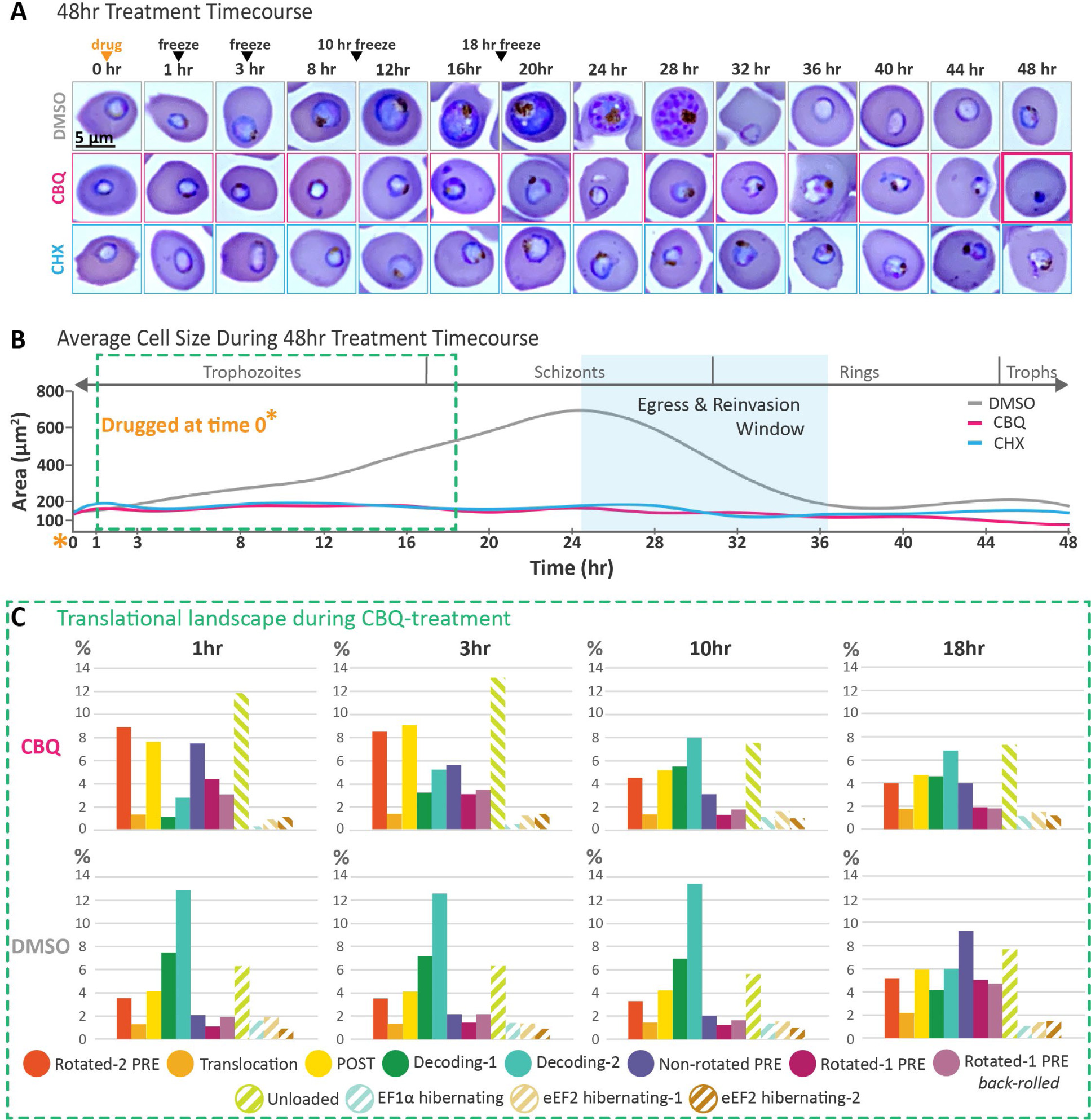
Phenotypic changes and shifts in translational landscape induced by CBQ-treatment over 48 hours. **A**, Giemsa-stained blood smears of highly synchronized early trophozoite-stage *P. falciparum* parasites, taken at 0, 1, and 3 hours after addition of DMSO, CBQ, or CHX, and every 4^th^ hour after, out to 48 hours. Three biological replicates were performed, with **n=25-81** for each time point in each replicate. One representative replicate is shown. Time of drug addition and time points corresponding to those used in our *in situ* cryoET studies are indicated with an orange and black arrowheads, respectively. Box with bold pink outline indicates a pyknotic cell. **B**, Plot comparing average parasite size at each timepoint for the DMSO-, CBQ-, and CHX-treated parasites. Time of drug addition is indicated with an orangeasterisk. Decrease in average parasite size in CBQ- and CHX-treated parasites starting at ∼20 hrs post-treatment is a consequence of increasing percentages of pyknotic parasites (small cell size). Average parasite size in DMSO-treated control parasites decreases during the window for merozoite egress and reinvasion into new erythrocytes, shaded in light blue, as schizonts (large parasite cell size) egress and reinvade, becoming newly invaded ring-stage parasites (small parasite cell size). **C**, Bar graphs showing the translational landscape comprised of relative distribution of identified translation intermediates and ribosomal states in CBQ-treated and DMSO-control parasites. Percentages are based on total amount of identified ribosomes after particle curation.

Our *in situ* cryoET datasets of drug-treated and DMSO-control parasites harvested at 1, 3, 10 and 18 hours post-treatment were collected based on these results. Across the 10 datasets encompassing four timepoints, we collected a total of 920 tomograms from 77 lamella containing 132 cells (Table S1). Examination of the relative distributions of *Pf*80S particles between the ribosomal states described above revealed shifts in the translational landscape in response to CBQ-treatment (Figure 4C).

At 1 hr post-treatment, CBQ-treated parasites exhibit a dramatic accumulation of ribosomes in states directly preceding or following the binding or dissociation of an elongation factor (either eEF2 or eEF1α) at the ribosomal GTPase site. This accumulation subsides with prolonged CBQ-exposure. Additionally, we observe a dramatic depletion of the eEF1α-bound decoding states at 1hr post-CBQ treatment compared with DMSO-control parasites, which recovers over time, but never returns to the levels exhibited in the control parasites. Interestingly, we observe that the translational landscape stays quite uniform in DMSO-control parasites in the first 10 hours, followed by a marked shift between the 10 and 18 hour timepoints, correlating with a shift in the translational activity of the parasites as they transition from trophozoites to schizonts (Figure 1A, 4A,B).

### Proteins involved in ribogenesis are enriched in parasites after prolonged CBQ exposure

To understand how inhibiting translation affects the *P. falciparum* proteome in the highly translationally active trophozoite stage, we performed quantitative proteomics on drug-treated early-trophozoite stage parasites harvested at 1, 3, 10, and 18 hours post-treatment. A comparison of bulk protein levels between CBQ-treated and DMSO-control parasites revealed reduced protein synthesis in drug-treated parasites starting as early as 1 hour post-treatment (Figure 5A, S5A), consistent with previous studies^37^. Bulk protein content is dramatically reduced by 10 and 18 hours post-treatment (Figure 5A, S5A).

**Figure 5.**
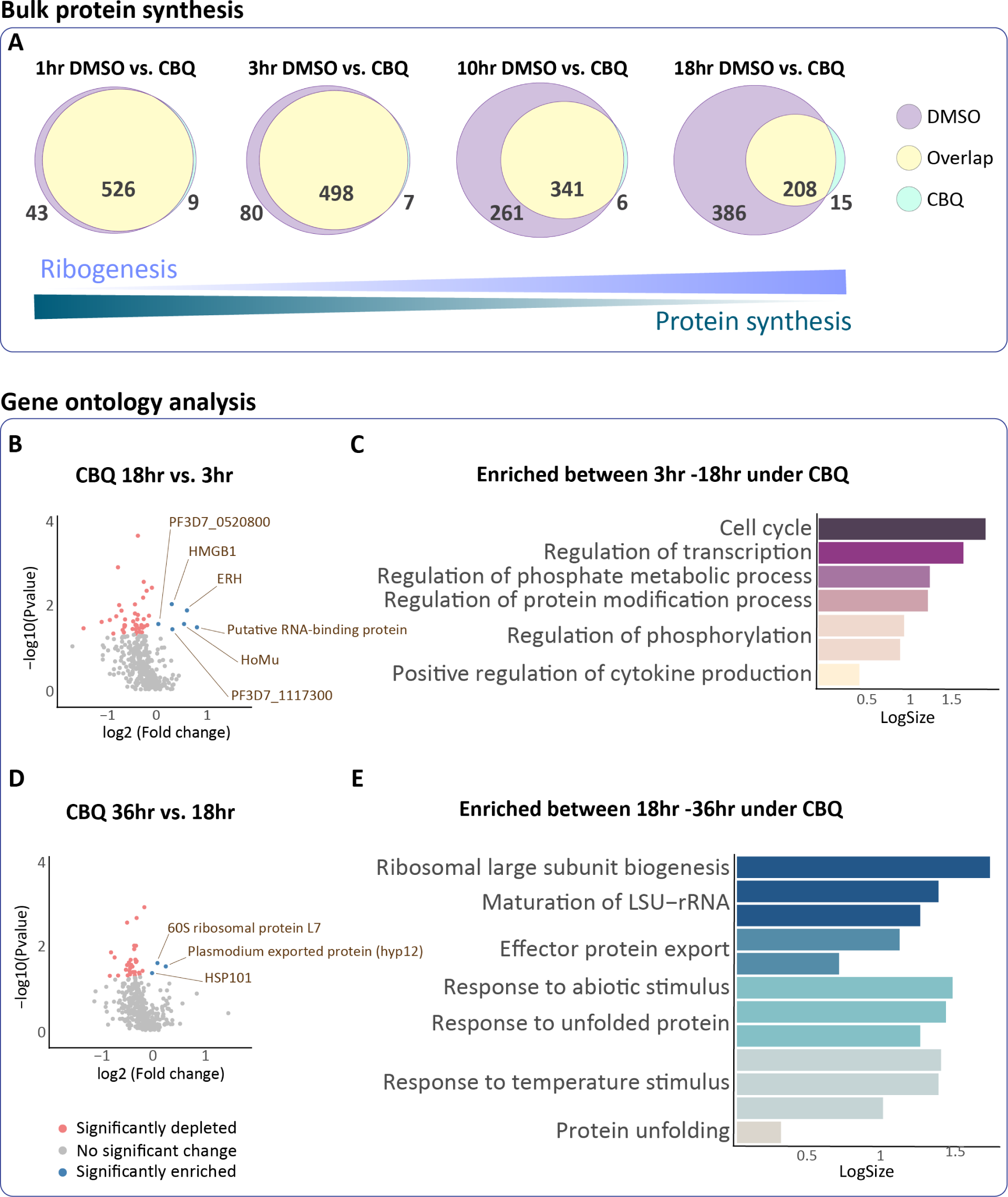
Quantitative proteomics reveals consequences of translation inhibition in *P. falciparum*. **A**, Venn diagrams showing differentially expressed proteins between DMSO-and CBQ-treated parasites at 1, 3, 10 and 18 hours post-treatment. **B,D,** Volcano plots showing differentially expressed proteins between different timepoints in CBQ-treated parasites comparing 18 hours to 3 hours **(B)** and 36 hours to 18 hours **(D)**. **C,E,** Revigo-generated GO enrichment analysis of enriched proteins in CBQ-treated parasites at 18 hours compared to 3 hours **(C)** and 36 hours compared to 18-hours **(E)**. For all volcano plots, the mean average of each protein from three replicates (n=3) is shown. Y-axes show log10 p-value and x-axes show log2 fold change using nested study design. Dark coral and blue circles indicate proteins with significant change. Grey circles indicate proteins with no significant (p>0.05) change in expression. For GO enrichment analysis graphs, bars are colored by Revigo-designated cluster. Y-axes are log10 of number of annotated GO term in *P. falciparum*. Each bar in the plot represents a sub-cluster.

To explore the effects of prolonged exposure on the proteome, we performed additional quantitative proteomics on CBQ-treated parasites harvested at 3, 18 and 36 hours post-treatment (Figure 5B-E). In this time frame, the DMSO-treated parasites have progressed normally through the schizont stage, replicated, egressed and reinvaded by ∼30 hours post-treatment, and their proteomes have cycled through three distinct blocs of gene expression profiles^5^ (Figure 4A). Consequently, we used the proteome of the 3 hour CBQ-treated parasites as a baseline for determining protein enrichment or turnover at the 18 and 36 hour timepoints. We observed enrichment of a small number of proteins in CBQ-treated parasites across both the 3-18 and 18-36 hour intervals, despite an overall depletion of proteins (Figure 5B,D). We then performed gene ontology enrichment (GO) analysis on these differentially regulated proteins in PlasmoDB^45^(Figure 5C,E, S5B-C), and were interested to note an enrichment of proteins involved in ribosome biogenesis and rRNA maturation between 18-36 hours (Figure 5E). This observation is further supported by a more granular comparison of the proteome in CBQ-treated parasites between 1-3 and 10-18 hours, which reveals a similar enrichment of proteins involved in ribogenesis across both time intervals, as well as an enrichment of translation-related proteins between 10-18 hours, beyond what would be expected based on the DMSO-treated control parasite data (Figure S5E-J). Curiously, we also observed enrichment of proteins involved in tRNA thio-modification (S^2^) (Figure S5J), which is known to be important for avoiding slowed translocation due to ribosome stalling in yeast^46^.

A similar analysis of CHX-treated positive control parasites across the same time intervals reveals an emphasis on translation and host hemoglobin metabolism (Figure S6A-G). These discrepancies in the way parasites respond to CBQ and CHX suggest differences in modes of cell death induced by these two drugs.

Meanwhile, analysis of global perturbations in the proteome under MMV019189-pressure revealed marginal bulk deviation from DMSO-treated control parasites at 1 and 3 hours (Figure S6H). GO analysis of specifically upregulated proteins in this time frame highlights prioritization of proteins involved in transcription and export of effector proteins into the host cell (Figure S6I-J). In direct contrast to CBQ-treated parasites (Figure S5H), proteins involved in cytoplasmic translation and amino acid metabolic processes, as well as regulation of proteasome assembly are depleted rather than enriched in MMV019189-treated parasites (Figure S6K). These processes collectively point to disruption of cytoplasmic translation and de-prioritization of hemoglobin uptake and amino acid biosynthesis in response to MMV019189.

### Visualizing *Pf*80S ribosomes in the native cellular context reveals evidence of increased ribogenesis with prolonged CBQ-exposure

To gain a comprehensive understanding of the effect of translation inhibition on *P. falciparum*, we integrated our local and global data by first mapping our subtomogram averaged *Pf*80S particles back into their originating tomograms (Figure 6-7). From this mapping back, membrane-bound ribosomes were identified and picked by hand, then validated by classification and refinement in Relion3.1 (Figure S4M-N). As seen in previous *in situ* studies^25,36^, we observed density corresponding to neighboring ribosomes in our unmasked *Pf*80S refinements, allowing us to measure a polysomal inter-ribosome distance of 15nm (Figure S4H-J). As such, we were able to locate tightly packed polysomes in our tomograms by identifying ribosome pairs where the distance between the mRNA entry and exit points fell within 15nm (Figure S4 K-L). To gain the full cellular context, we then mapped averaged central slices from tomograms collected on each cell back onto a lower-magnification image of the cell, then overlaid ultrastructural segmentations and mapped back ribosome reconstructions on top, yielding an integrated cell montage spanning multiple length-scales (Figure 6A, 7A).

**Figure 6.**
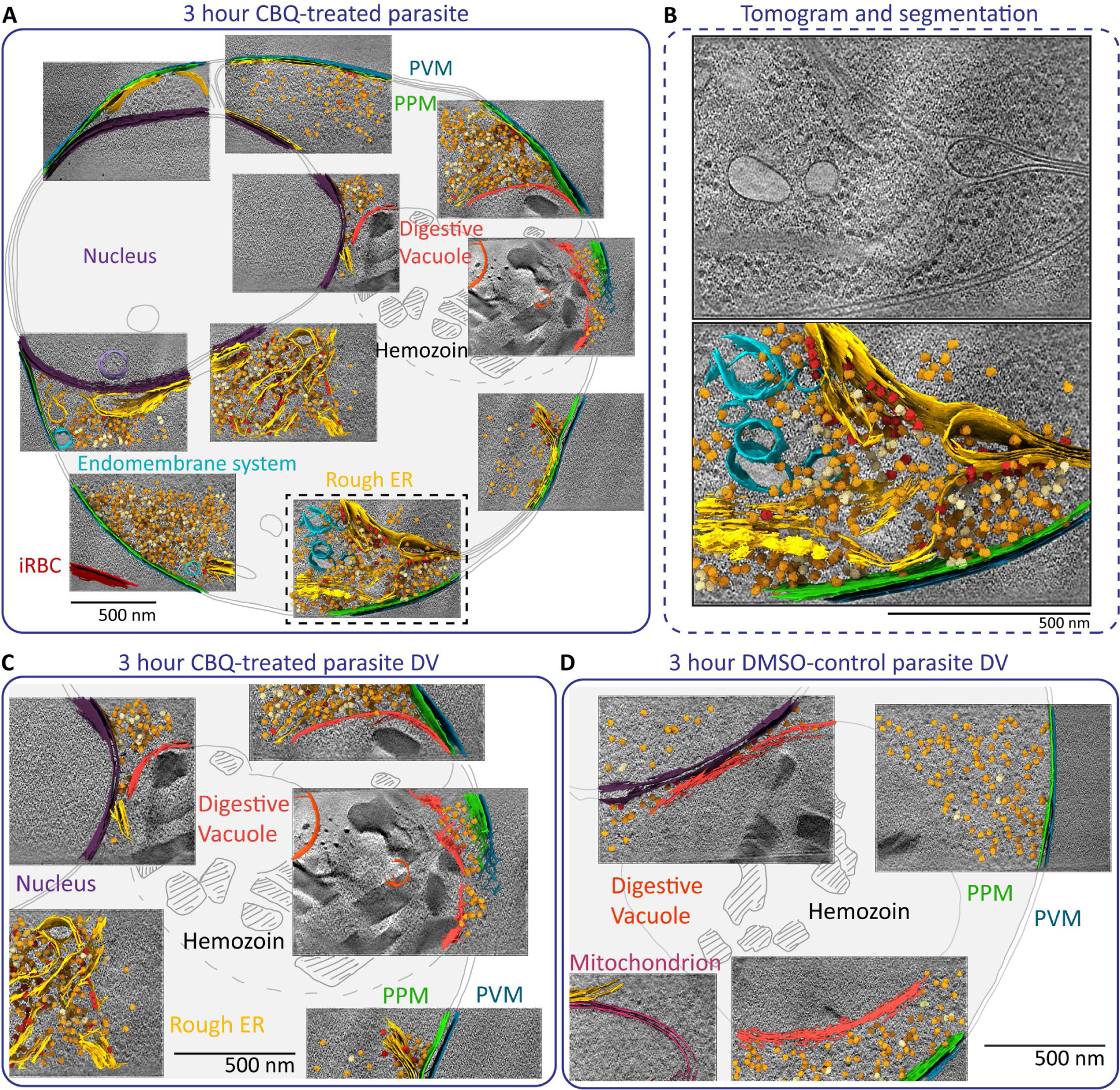
Integrated cell montages of CBQ- and DMSO-treated *P. falciparum* parasites at 3 hours post-treatment. **A,** Integrated cell montage of a CBQ-treated parasite frozen, milled and imaged at 3 hours post-treatment. Averaged central slices from tomograms collected on the cell are shown mapped onto a tracing of the cell, with ultrastructural segmentations and mapped back ribosome reconstructions overlaid on top. Segmented features are color coded as indicated by corresponding colored labels. Mapped back ribosomes are colored as follows: orange=monomers, pale-yellow=dimers and polysomes, red=membrane bound ribosomes. **B**, Detailed view of the tomogram indicated by dashed box in (**A**), shown alone (top) and overlaid with corresponding segmentation (bottom). Segmentation is colored as in (**A**). **C**, Detailed view of the averaged central slices overlaid with segmentations containing the DV of the cell shown in (**A**). Segmented features are color coded as in (**A**). **D**, Detailed view of the DV from a 3 hours DMSO-control parasite. Averaged central slices of tomograms containing the DV are overlaid on a tracing of the corresponding portion of the cell with segmentations. Segmented features are color coded as in (**A**). PVM, parasitophorous vacuolar membrane. PPM, parasite plasma membrane. EMS, endomembrane system. DV, digestive vacuole. Hz, hemozoin. ER, endoplasmic reticulum.

**Figure 7.**
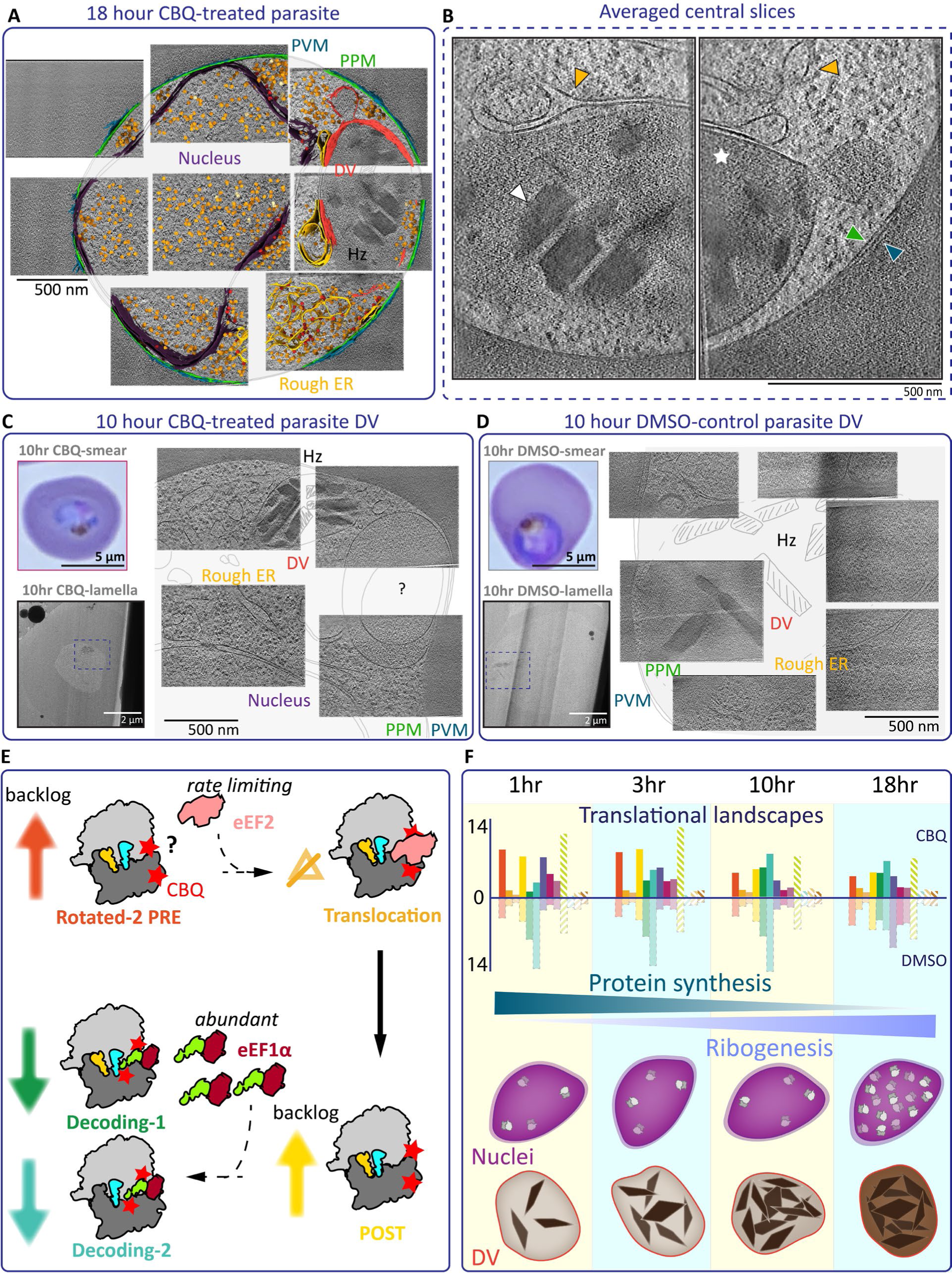
Integrated cell montages of CBQ-treated *P. falciparum* parasites at 18 and 10 hours post-treatment. **A**, Integrated cell montage of a CBQ-treated parasite frozen, milled and imaged at 18 hours post-treatment. Segmented features are color coded as in Fig 6. **B**, Detailed view of the averaged central slices containing the DV of the cell shown in (**A**). Colored arrowheads are color-coded as in (**A**). White arrowhead indicates a hemozoin crystal. White star indicates the DV lumen. **C**, Giemsa stain of a 10 hr CBQ-treated parasite. Lamella image of vitrified and milled 10 hr CBQ-treated parasite. Close-up view of the parasite in the lamella. Detailed view of the DV from the cell. Averaged central slices of tomograms containing the DV are overlaid on a tracing of the corresponding portion of the cell. **D**, Giemsa stain of a 10 hr DMSO-control parasite. Lamella image of vitrified and milled 10 hr DMSO-treated parasite. Close-up view of the parasite in the lamella. Detailed view of the DV from the cell. **E**, We propose that CBQ (red stars) perturbs eEF2 (salmon) and eEF1α-A-tRNA (dark red, light green) interaction with the *Pf*80S GTPase site, preventing smooth transition into the translocation and decoding states, respectively. Failure to progress into these states of translation elongation leads to an accumulation of of rotated-2 PRE (dark orange arrow) and POST (yellow) states (Fig 4**C**). **F**, Summary of CBQ effect on trophozoite stage parasite nucleus, digestive vacuole, gene expression pattern and translational landscape across 18 hr time course.

In line with our Giemsa-stained blood smears, CBQ-treated parasites in our integrated cell montages remain consistent in size and organellar organization across all timepoints (Figure 6A, 7A). Conversely, DMSO-control parasites increase dramatically in size and organellar content beyond the 3-hour timepoint (Figure 7D, S7C-E). By 18 hours post-treatment, the appearance of additional nuclei, sometimes bound by additional membranes, in DMSO-control parasites indicate the beginning of the transition from trophozoites to schizonts (Figure S7C-E). We observe membrane-bound ribosomes distributed on rough endoplasmic reticulum (ER) and nuclear envelope membranes (Figure 6A-C, 7A). The rough ER is abundant and distributed throughout the cell, and we find segments that are continuous with the outer leaflet of the nuclear envelope, as well as points that are in direct apposition with the parasite plasma membrane or DV (Figure 6A, C, 7A). Strikingly, the nuclei in our CBQ-treated parasites contain an unusual abundance of ribosomes in the nucleus at 18 hours post-treatment (Figure 7A), further substantiating our proteomic evidence (Figure 5E) that ribosome biogenesis is upregulated in response to prolonged CBQ-exposure.

### Digestive vacuole abnormalities suggest prolonged CBQ exposure disrupts hemoglobin metabolism

We observe prominent ultrastructural abnormalities in the parasite-specific digestive vacuole (DV) in CBQ-treated parasites that progress in severity with prolonged drug exposure (Figure 6A, C, 7A-C). The digestive vacuole is a unique organelle used by the parasite to recycle hemoglobin taken up from the host erythrocyte into amino acids and heme^47^. This key source of amino acids is essential for parasite growth and replication, but the released heme forms radicals that are toxic to the cell. To protect itself, the parasite biocrystallizes much of the released heme into inert hemozoin crystals inside the DV^48^. Despite this protective measure, the concentration of labile heme in the parasite cytosol (∼1.6mM) far exceeds normal cytosolic heme concentrations in other organisms (∼20-600nm)^49^. This may explain, in part, the unusually high background in our tomograms from milled intraerythrocytic parasites.

In DMSO-control parasites, we observe that hemozoin crystals are generally well-distributed inside the DV lumen, and the area in the DV lumen surrounding the crystals is noticeably less electron-dense than the erythrocyte cytosol (Figure 6D, 7D). In CBQ-treated parasites, compaction of the DV lumen around the hemozoin crystals can be seen as early as 3 hours post-treatment (Figure 6A, C). Crystals pack more densely and free space in the DV lumen decreases even further at 10 hours post-treatment (Figure 7C). At 18 hours post-treatment, the DV lumen looks almost as electron dense as the hemoglobin-rich erythrocyte cytosol (Figure 7A-B), much more electron dense than the surrounding parasite cytosol. This closely resembles phenotypes induced by the inhibition of DV proteases, suggesting the hemoglobin is no longer being broken down properly inside the DV in our CBQ-treated parasites^50^ (Figure 7A-B).

## DISCUSSION

The *Plasmodium*-specific translation inhibitor CBQ has been shown to inhibit protein synthesis in malaria parasites^37^. *P. falciparum* translation elongation factor 2 (*Pf*eEF2) is thought to be the target, as mutations in *Pf*eEF2 are found in all recrudescent parasites arising under CBQ pressure^37^. However, the mode of action of CBQ remains unclear.

We propose that CBQ disrupts protein synthesis by directly or indirectly perturbing interaction of both eEF2 and eEF1α with the *Pf*80S ribosome (Figure 7E). In support of this hypothesis, in our comparison of the translational landscape between CBQ-treated and DMSO-control parasites we observe a relative increase in all non-GTPase-bound states in response to CBQ exposure, suggesting ribosomes are accumulating in these states due to a backlog caused by less efficient progression through the GTPase-bound states. Specifically, we observe a marked decrease in the eEF1α-bound decoding states in response to CBQ-exposure, indicating that CBQ is preventing eEF1α binding to the ribosome. Furthermore, we observe an accumulation of ribosomes in the POST state directly preceding the decoding states, further supporting that transition to the eEF1α-bound decoding-1 state is inhibited by CBQ.

Intriguingly, we see a similar accumulation of ribosomes in the rotated-2 PRE state directly preceding the translocation (eEF2-bound) state in response to CBQ-treatment, indicating that progression to the translocation state is negatively impacted by the presence of CBQ. However, in contrast to the dramatic effect of CBQ on the decoding states, the relative abundance of the translocation state remains relatively unchanged in response to CBQ-treatment. Since levels of ribosomes in the translocation state are not reduced, a mechanism by which CBQ prevents eEF2 binding can be excluded. This leaves the possibility of CBQ affecting eEF2 movement or dissociation. The fact that the relative abundance of ribosomes in the translocation state remains fixed across all timepoints in both DMSO-control and CBQ-treated parasites suggests the availability of activated eEF2 is rate-limiting and the maximum number of activated eEF2 molecules are always engaged. Although *in situ* cryoET enables us to visualize these events in incredible detail, these snapshots in time do not provide insight into the effect of CBQ on the kinetics of eEF2 movement or dissociation, which requires further investigation.

While the resolution of each of our states is insufficient to allow us to see where CBQ binds, we imagine a model in which CBQ binds somewhere in the GTPase site and directly modulates the binding of factors at that site, given that we observe accumulation of ribosomes in the state directly preceding each elongation factor binding event in response to CBQ-treatment. It is also possible that, similar to other translation inhibitors like cycloheximide^43,51^ and emetine^10^, CBQ binds at the interface between the large and small subunits near the E site in a way that impedes rotations of the 40S required for transition between different states. Our data advances understanding of how CBQ impedes malarial translation elongation, informing further study and therapeutic development.

Taken together, our work transforms the current understanding of translation elongation in *P. falciparum*. By showing that *Pf*RACK1 is associated with all ribosomal states observed *in situ*, we have resolved the controversy surrounding *Pf*RACK1 and the *Pf*80S ribosome. Our discovery of a bifurcated pathway during the transition from the decoding to peptidyl-transfer states in translation elongation raises questions about whether this represents an evolutionary adaptation specific to *P. falciparum* or an as-yet-undescribed general phenomenon. Our time-resolved, multi-scale interdisciplinary approach reveals drug-induced perturbations in the translational landscape, proteome and parasite ultrastructure, despite the apparent inactivity reflected in blood smears (Figure 7F). Of note, the increase in ribosome biogenesis as revealed by our proteomics and molecular and ultrastructural cryoET data is an intriguing stress response inviting further investigation (Figure 7F). Our work informs the targeted development of new therapies and prompts further study of protein synthesis in malaria parasites as well as a broader exploration of how the utilization of translation intermediates varies across organisms.

## ACKNOWLEDGMENTS

We acknowledge support from the National Institutes of Health (DP5OD029613, CMH), the Columbia Precision Medicine Pilot Initiative (CMH), the Swiss National Science Foundation (P2BSP3_200205), and the Human Frontiers Science Program (LT000452/2021-L). We thank the Weill Cornell Medicine Proteomics and Metabolomics Core Facility and Indiana University School of Medicine Center for Proteome Analysis for assistance in mass spectrometry. All cryoFIB-milling was performed on a TFS Aquilos cryoFIB-SEM owned by the Fitzpatrick Lab in the Zuckerman Mind Brain and Behavior Institute at Columbia University. CryoET data was collected on Titan Krios instruments in the Columbia Electron Microscopy Center, housed either in the New York Structural Biology Center or the Zuckerman Mind, Brain and Behavior Institute at Columbia University, and on the Titan Krios instrument at the University of Colorado Boulder Biochemistry Department’s Krios Electron Microscopy facility (BioKEM). We thank [Daniel Goldberg, Rachel Green, Kirk Deitsch, David Fidock, Alan Brown, Björn Kafsack, Ahmad Jomaa, Anthony Fitzpatrick, Israel Fernandez, and Hamna Shahnawaz] for helpful discussions and suggestions.

## AUTHOR CONTRIBUTIONS

CMH, LA and DWC initiated the project. CMH, WC and MTH designed drug treatment time-course experiments and strategy for preparing parasite samples for proteomics. WC performed drug treatment time-course experiments and interpreted the results. MTH prepared cultured parasites for proteomic studies. CMH and MTH interpreted the proteomics results. HK prepared crudely enriched ring-stage lysates. GZ performed proteomics data collection and analysis. EHD performed proteomics data analysis. CLL helped with all parasite culture. DWC, WC and MTH cultured, froze, and screened parasites for the untreated trophozoite and schizont, drug- and DMSO-treated trophozoite, and merozoite datasets, respectively. CMH developed protocols for cryoFIB-milling of frozen parasites and tilt series collection on the resulting lamella. CMH and WC performed cryoFIB-milling for all datasets. MTH helped perform cryoFIB-milling for merozoite and drug- and DMSO-treated trophozoite datasets. CMH, LA and WC collected tilt series for all datasets. CMH, LA and WC developed the tomogram reconstruction pipeline. LA, WC and XZ manually picked and curated particles for subtomogram averaging. CMH and LA developed and performed subtomogram averaging and interpreted the resulting structures and their distributions. LA built and refined the atomic model. CMH, LA, LH, EL, and AN adapted published code for mapping back of subtomogram averaged ribosomes into tomograms and identifying polysomes in tomograms. WC performed tomogram denoising and segmentation with help from XZ. CMH, LA, WC, and MTH wrote the paper. All authors discussed and commented on the manuscript. CMH supervised all aspects of the project.

## Methods

### Parasite Culture

Asexual *P. falciparum* parasites were cultured as in Drew et al, 2008 [^52^] in O+ or A+ erythrocytes (New York Blood Center), at 37°C under 5% O_2_, 5% CO_2_, 90% N_2_ in complete RPMI (cRPMI): RPMI 1640 medium supplemented with 0.05 mg/mL hypoxanthine, 0.06 mg/mL sodium hydroxide, 0.8 mg/L thymidine, 0.04 mg/mL sodium pyruvate, 2.25 mg/mL sodium bicarbonate, 5.9 mg/mL HEPES, 0.67 mg/mL glucose, 0.01 mg/mL Gentamycin, and 0.5 % Albumax II (Gibco), at a hematocrit of 2%. Unless otherwise specified, reagents were purchased from Sigma.

### Crude fractionation of ring stage parasites

Tightly synchronized parasites were harvested at the ring stage at 8-15% parasitemia by pelleting and then re-suspending in AIM buffer (KCl 120 mM, NaCl 20mM, Glucose 20mM, HEPES 6mM, MOPS 6mM, MgCl2 1mM, EGTA 0.1mM, pH 7.0) and equal volume of AIM buffer supplemented with 0.1% saponin. After washing the pellet 3 times with AIM buffer, pellet was resuspended in MESH buffer (Manitol 225mM, Sucrose 75 mM, MgCl2 4.3mM, Tris 10mM, EGTA 0.25 mM, HEPES 15 mM, pH 7.6) supplemented with 1mM PMSF and 1µL/mL fungal protease inhibitor cocktail. Parasites were slowly disrupted by N2 cavitation (4639 Cell Disruption Bomb, Parr, USA) at 1,000 psi at 4°C, then spun down at 900 g for 5 min to separate solid from soluble fraction. The solid phase containing nucleus and large debris was removed. The cloudy supernatant was passed through a magnetic column (Miltenyi Biotec) to remove hemozoin. The elute was pelleted by centrifugation at 20,000 g for 30 min at 4°C. Pelleted fractions were flash frozen in liquid nitrogen and stored at -80°C. Samples were thawed and exchanged into 20mM HEPES pH 7.5, 150mM NaCl, 5mM MgCl2 directly before freezing grids for *ex vivo* cryoET.

### Parasite Enrichment

To enrich merozoites, Sorbitol (Fisher Scientific, AA3640436) synchronized NF54 *P. falciparum* parasites were allowed to progress to the schizont stage, purified using a 65% Percoll (Cytiva) cushion and then treated with 10 µM E-64 (Sigma, E3132) for 4-8 hours until mature merozoites had formed. Merozoites were then mechanically released from the host erythrocyte by passage through a 1.2 µm syringe filter (Pall Life Sciences, 4656), pelleted at 2000 x g for 5 min, and treated with 16 µM DAPI (Thermo Scientific, 62249) prepared in cRPMI for 5 min at 37°C. The DAPI was then washed out and the cells resuspended in cRPMI.

To enrich trophozoites, Sorbitol synchronized Exp2-mNG parasites^50,53,54,50,53^ were allowed to progress to the trophozoite stage. Drug treated trophozoites were treated with the appropriate drug prepared in cRPMI or with cRPMI containing DMSO-vehicle control. Drugs used: 20 nM CBQ (ApexBio, A8711) prepared in DMSO and 15 µM MMV019189 prepared in DMSO. Parasite-infected red blood cells (iRBCs) were harvested after 1, 3, 10 and 18 hours of treatment. For each timepoint, an aliquot of purified trophozoites were placed back into culture to monitor parasite development over the following 48 hours of drug exposure. Blood smears of all parasites were stained using Hema3 Fixative and Solutions (Fisher Healthcare) and imaged at 100x magnification using an Echo Revolve R4 microscope at each timepoint.

### Giemsa-stained 48hr Drug Treatment Timecourse

Parasites at trophozoite stage were synchronized and purified by Percoll followed by incubation in 0.7% gelatin at 37°C for 1 hour. Purified trophozoites were resuspended in RPMI with 20 nM CBQ (DMSO being the control). Purified trophozoites were resuspended in 1 mL RPMI at 5% hematocrit into a 24-well plate (Corning). A small aliquot of culture was collected for blood smears every 4 hours over the next 48hr.

### Proteomics

Synchronized EXP2-mNG parasites were expanded to 1.3 L and treated with 20nM CBQ, 15 µM MMV019189, 4 µM Cycloheximide, or DMSO in cRPMI and incubated at 37°C. Treated parasites were harvested by washing twice with 1xPBS and flash frozen and stored at -80°C. TMT based quantitative proteomics was performed at Weill Cornell Medicine Proteomics and Metabolomics Core Facility. For all experiments, cells were grown and treated in biological triplicate. For the first round of proteomic analysis, DMSO-, CBQ- and MMV019189-treated infected RBC pellets were harvested at 1hr and 3hr post-treatment. For the second dataset, DMSO-, CBQ-, and CHX-treated pellets were collected at 10hr and 18hr post-drug treatment. For the third dataset, CBQ-, and CHX-treated pellets were harvested at 3hr, 18hr and 36hr post-treatment. For the third dataset, a concurrently cultured DMSO-treated sample was smeared at 3hr, 18hr and 36hr post-treatment to confirm normal parasite development.

Pellets were provided to the facility where proteins were extracted, reduced, alkylated and digested with trypsin. Desalted peptides (50 ug) were labeled by 18-plex TMTpro (Thermo Fisher Scientific) of which a small aliquot was mixed and analyzed by LC-MS to determine labeling efficiency and necessary ratio mixing. All samples were mixed at equal ratios and fractionated by offline RPLC into 12 fractions. 5% of each fraction was run by LC-MS for global proteomics and the remaining 95% was enriched for phosphopeptides using TiO2 beads (GL Sciences). An EASY-nLC 1200 coupled on-line to a Fusion Lumos mass spectrometer (Thermo Fisher Scientific) was used for LC-MS. Buffer A (0.1% FA in water) and buffer B (0.1% FA in 80% ACN) were used as mobile phases for gradient separation. A 75 µm x 15 cm chromatography column (ReproSil-Pur C18-AQ, 3 µm, Dr. Maisch GmbH, German) was packed in-house for peptide separation. Peptides were separated with a gradient of 10–40% buffer B over 110 min, 40%-80% B over 10 min at a flow rate of 300 nL/min. The Fusion Lumos mass spectrometer was operated in data dependent mode. Full MS scans were acquired in the Orbitrap mass analyzer over a range of 400-1500 m/z with resolution 60,000 at m/z 200. The top 15 most abundant precursors with charge states between 2 and 6 were selected with an isolation window of 0.7 Thomson by the quadrupole and fragmented by higher-energy collisional dissociation with normalized collision energy of 40. MS/MS scans were acquired in the Orbitrap mass analyzer with resolution 30,000 at m/z 200. The automatic gain control target value was 1e6 for full scans and 5e4 for MS/MS scans respectively, and the maximum ion injection time was 100 ms for MS scans and 54 ms for MS/MS scans.

Data analysis was performed at the Center for Proteome Analysis at IUSM using Proteome Discoverer 2.5 (Thermo Fisher Scientific). The raw data was searched against *Homo sapiens* (downloaded 05/13/22, 20292 sequences), *Plasmodium falciparum* 3D7 (downloaded 04/06/21, 5381 sequences) and common laboratory contaminants (73 sequences) protein databases using Sequest HT. Full trypsin digestion with a maximum of 3 missed cleavages, a precursor mass tolerance of 10 ppm and a fragment mass tolerance of 0.02 Da were used. Static modification of carbamidomethyl C was utilized. Dynamic peptide modifications were set at a maximum of 3 per peptide including oxidation of M; phosphorylation of S, T, and Y; Deamidation of N and Q; and TMTpro of K and peptide N-terminus; and Acetyl-Met-loss or Met-loss plus acetyl on protein N-terminus. Percolator FDR cutoffs of 0.01% strict and 0.05% relaxed at the PSM level were employed. The IMP-ptmRS node was used for phosphosite localization confidence scoring. At the consensus level, unique and razor peptides were used for quantification with reporter abundance based on S/N, isobaric quantification corrections applied, a co-isolation cutoff of 30% and a S/N threshold cutoff of 6. Normalization was done using total peptide amount with no scaling. Results were filtered for *Plasmodium falciparum* proteins and phosphopeptides and exported to excel (Microsoft).

Relative quantitation of protein and phosphopeptide intensities were log transformed and normalized using median intensity value per sample. One way ANOVA and pairwise comparison was used to compare each condition. Volcano plots showing differential expression of proteins were generated using ggplot in R Studio. Gene Ontology analysis of differentially expressed proteins was done with the GO Enrichment tool in PlasmoDB (https://plasmodb.org) focused on biological processes with P-value cut-off of 0.05. Generated GO terms were imported into the REVIGO (Reduce and Visualize Gene Ontology) web server and analyzed for clustered visualization.

### Sample freezing and cryoCLEM

For *in situ* cryoET, 3.5 µL of parasite-infected RBCs in cRPMI were applied to glow-discharged R2/2, 200 mesh Au, or Cu, carbon or SiO_2_ Quantifoil EM grids, followed by manual back-blotting with Whatman # 1 filter paper and plunge freezing in liquid ethane using a custom-built manual plunger. Frozen grids were clipped into AutoGrids (Thermo Scientific) and assessed for ice thickness and cell density using a Cryo Correlative Light and Electron Microscope (cryoCLEM) (Leica Microsystems). For *ex vivo* cryoET, crudely fractionated lysate from untreated *P. falciparum* ring stage parasites were plunge frozen in liquid ethane in a Vitrobot MarkIV semi-automated plunge freezer (Thermo Scientific) on glow-discharged R2/2, 200 mesh, Au, carbon Quantifoil EM grids (Quantifoil Micro Tools).

### Focused Ion Beam-Scanning Electron Microscopy

Grids were loaded into an Aquilos cryo focused ion beam scanning electron microscope (Thermo Scientific) and lamellae were created as previously described^55^. In brief, eucentric height was determined for suitable target sites, after which the grid was sputter coated with platinum metal, then coated with trimethyl(methylcyclopentadienyl)platinum(IV) using the onboard gas injection system (GIS). Tension release trenches were milled at an angle of 40°, after which the milling process was performed at angles of 7°-8° at 30 kV, using currents of 0.5 nA, 0.3 nA., 0.1 nA, 50 pA and 10 pA for the focused ion beam. Final lamellae were polished to a thickness of 150-200 nm.

### Data Collection

Data collection was performed on three high end cryoTEM microscopes operating at 300kV: Two Titan Krios (Thermo Fisher) with K3 cameras and Gatan Imaging Filter systems (Gatan) and one Titan Krios with Falcon IV (Thermo Fisher) and Selectris energy filter (Thermo Fisher) (Table S1). All data was collected in super-resolution mode at a magnification of 42,000x (physical pixel size of 2.094Å/px) or 53,000x corresponding to a physical pixel size of, 1.66, 1.699 or 1.6 Å/px, depending on the microscope (Table S1). SerialEM (for K3 camera)^56^ or EPU Tomo (for Falcon IV camera) (Thermo Scientific) acquisition softwares were used to set up and collect tilt series with a total dose of 120e-/Å^2^, over 41 tilts comprising 10 frames per tilt. Each tilt series was collected over a range of 120° in 3° increments following a dose-symmetric scheme, starting at a pre-tilt angle defined by the milling angle (7°-8°).

### CryoET Data Processing and Analysis

Collected tilt series (Table S1) were pre-processed using Warp 1.0.9 [^57^] including 2D CTF correction, Motion Correction and creation of tilt stacks. Tomogram alignment and reconstruction were done without pretilt correction in AreTomo^58^ and denoised using Topaz 3D denoise^59^.

#### Subtomogram Averaging (STA)

The crYOLO semi-automated particle picker^60^ was manually trained on each dataset and the resulting model applied to all tomograms of the same dataset (Table S1). The resulting particle picks were manually curated in EMAN2 e2spt_boxer.py^61^ to exclude false positives. Subtomograms were extracted in Warp 1.0.9 using alignment files from AreTomo and averaged and refined in RELION 3.1 [^62^]. Resulting reconstructions and particle orientations were imported into M 1.0.9 [^63^] to sequentially refine particle pose, image and volume warp and defocus parameters. Improved particles were re-extracted and again refined in RELION. This scheme was continued until the consensus refinement at 2.094 Å/px reached Nyquist of 4.2 Å and was used for 3D focused classifications (Figure S2, S4A). Particles from ring, trophozoite, schizont and merozoite datasets collected at 1.7 Å/px were extracted separately to full pixel size and refined again in RELION3.1 for final consensus at an overall resolution of 4.1Å (Figure 2A-C, S1A-C, Table S1, S2).

#### Focused 3D classification

To improve *Pf*RACK1 density, focused classification of the 40S body was performed in RELION3.1, leading to separation of rotated (rt*Pf*80S) and non-rotated (nrt*Pf*80S) *Pf*80S (Fig. 2D-E, Fig. S2B). The nrt80S particles were further refined by centering on the head region, improving local map quality and resulting in a final 4.3 Å reconstruction (Figure 2D-M, S2B).

To separate the individual translation intermediates, several rounds of focused classification were performed on all particles in the refined consensus map at 2.094Å/px (Fig. S2A, S4A). Classifications were performed in three steps. A first classification was performed to separate particles with and without density in the GTPase site using a mask around the GTPase site (EF mask). In a second step, particles with and without GTPase site density are both separately subjected to another round of classification, separating out the various elongation factor-bound or unclear GTPase site density states (eEF1α and eEF2) for particles containing density in the GTPase site, or separating out particles based on occupancy of the A or P tRNA sites for particles with no density in the GTPase site. In some cases, a mask around the A site tRNA (A-tRNA mask) was used to separate out the different elongation factors. In a third step, a mask around the P* and partial E site is used to separate out P* and E occupancies (P*E mask). Sometimes a fourth classification was necessary using the above-mentioned masks or a mask covering the GTPase, A, P and E sites (full mask) to get final classes. A full mask of the GTPase and tRNA binding sites was generated based on the refined consensus map in Xmipp. The GTPase site, A-tRNA site, and P*E site masks were generated from the full mask by deleting unwanted regions using the volume-erase function in ChimeraX. Soft masks were then created from these in RELION3.1_mask_create, using a 3-pixel binary mask extension and 4 pixel soft edge.

#### Calculating distribution of particles between ribosomal states, by dataset

All particles in each final classified state were refined in RELION3.1 to yield a final consensus reconstruction of each state. Details of the reconstructions of the individual states can be found in Table S2. To determine the distributions of particles between states for each dataset, we took the final RELION Refine3D star file for each individual state and wrote out a separate star file for each dataset, containing only the subset of particles originating from the corresponding dataset. This yielded the exact number of particles in each state from each dataset, from which we were then able to calculate the distributions of ribosomes in each state for each dataset, as a percentage of the total number of ribosomes in each dataset.

#### Segmentation

Segmentations were generated in a semi-automated manner using EMAN2. Briefly, models were trained on manual membrane annotations for each tomogram, then applied to the tomogram and then manually separated and colored for display and figure making in UCSF ChimeraX^64^.

### Atomic Model building

Models for *Pf*RACK1, *Pf*eEF1α_AF_ and *Pf*eEF2_AF_ were downloaded from the AlphaFold^19^ website under accession codes: AF-Q8IBA0-F1 (*Pf*RACK1_AF_), AF-QI80P6 (*Pf*eEF1α_AF_) and AF-Q9NDT2-F1 (*Pf*eEF2_AF_). SWISSMODEL^31–35^ was used to generate alternative models for *Pf*eEF1α and *Pf*eEF2 based on human eEF1α (PDBID: 8g6j) and human eEF2 (PDBID: 6z6m). In order to assess translational states and confirm binding positions, previously published models or maps of full 80S ribosomes were fitted into our *Pf*80S maps. The PDB model 3jbp was used as a basis for our new model. The model was ridged body fitted into the highest resolution consensus structure using ChimeraX. The *Pf*RACK1_AF_ model was placed based on h80S (6z6m) in ChimeraX and then added to the model in COOT^65^. Refinement and subsequent validation of the new model was done using the Phenix software suite^66^ (Table S3). Refinement strategy included minimization_global, local_grid_search, occupancy, adp and hqh_flips for 100 iterations and 5 macro cycles using secondary structure and NCS constraints or default settings.

### Computational polysome analysis

Polysomes were computationally identified using the relative distances between the entry and exit sites of all ribosomes in a tomogram (Figure S4H-J). First, exit and entry sites were defined as x-, y-, z-coordinates in the consensus reconstruction, relative to the point of origin in ChimeraX (Figure S4H). Then, the maximum allowed distance between exit of leading- and entry of following-ribosome in a *P. falciparum* polysome was defined by simulating placement of two ribosomes in a polysome based on low resolution densities for neighboring ribosomes in our reconstructions (Fig S4I), and confirmed with previously performed polysome analysis^15^. Measured distances between the simulated polysome was 138 Å, based on this a threshold of 150 Å was set (Figure S4J). The determined exit and entry site coordinates were then applied to the refined rotation angles and center of mass shifts for each particle, enabling calculation of the specific positions of entry and exit site of all individual ribosomes comprising the consensus reconstruction.

Using the newly generated coordinates, we identify the leading and following ribosome (Fig S4I) for each ribosome pair (i,j) by recording the distance between ribosome i’s entry from ribosome j’s exit at matrix index (i,j) and vice versa at matrix index (j,i). Between these two matrix indices, only the smaller index (shorter distance) is considered and the larger one is replaced with an arbitrarily large number. Next, we applied the defined distance threshold of 150 Å to this filtered matrix, setting all values above this threshold to 0. The nonzero entries of the new matrix are used to add ribosomes to disjoint sets, each set representing either a monosome (1 element), a dimer (2 elements) or a polysomes (>2 elements)(Figure S4K,L).

### Mapping back of ribosomes

The relionsubtomo2ChimeraX python script (DOI 10.5281/zenodo.6820119) was adapted to include polysome information. The script produces a .cxc file, containing information on ribosome poses in the original tomogram based on RELION star file input and enables positioning of reconstructions in the tomographic volume by ChimeraX. Based on results obtained from the polysome analysis (see previous section), the code assigns different colors to monomers (orange), dimers and polychains (butter yellow). The mRNA entry and exit sites for each ribosome can be displayed by green and yellow markers, respectively (Figure S4H).

### Figures and plots

All figures were generated in Adobe Illustrator and graphs plotted in Microsoft Excel or R studio. ChimeraX was used to visualize STA reconstructions and segmentations. Central slice images were generated in IMOD. Lamella montages were compiled in Adobe Photoshop.

---

**Table 2:** Final reconstructions of translation intermediates.

|  | Consensus | Rack1 | Rotated-2 PRE | Translocation | POST | Decoding-1 | Decoding-2 | Non-rotated PRE |
| --- | --- | --- | --- | --- | --- | --- | --- | --- |
| **Data Processing** |  |  |  |  |  |  |  |  |
| EMD | 41485 | 41494 | 41490 | 41491 | 41492 | 41486 | 41487 | 41488 |
| Symmetry imposed | C1 | C1 | C1 | C1 | C1 | C1 | C1 | C1 |
| Final particle images (#) | 120'226 | 63'941 | 16'495 | 10'283 | 20'415 | 13'275 | 13'542 | 14'578 |
| Pixel size (Å/px) | 1.69 | 2.094 | 2.094 | 2.094 | 2.094 | 2.094 | 2.094 | 2.094 |
| Map resolution (Å) FSC threshold 0.143 | 4.1 | 4.3 | 6.0 | 5.9 | 4.7 | 6.7 | 7.7 | 5.3 |
| Map resolution range (Å) | 3.4 - 10 | 4.2-11.2 | 4.5-14.5 | 4.4-13.3 | 4.2-10.1 | 4.6-12.6 | 6.1-14.3 | 4.2-11.5 |

|  | Rotated-1 PRE | Rotated-1 PRE<br>*back-rolled* | unloaded | eEF2, P? | eEF1α<br>hibernating | eEF2<br>hibernating-1 | eEF2<br>hibernating-2 |
| --- | --- | --- | --- | --- | --- | --- | --- |
| **Data Processing** |  |  |  |  |  |  |  |
| EMD | 41489 | XXXX | 41493 | XXXX | XXXX | XXXX | XXXX |
| Symmetry imposed | C1 | C1 | C1 | C1 | C1 | C1 | C1 |
| Final particle images (#) | 7'102 | 3'281 | 29'301 | 1'097 | 1'376 | 1'814 | 1'397 |
| Pixel size (Å/px) | 2.094 | 2.094 | 2.094 | 2.094 | 2.094 | 2.094 | 2.094 |
| Map resolution (Å) FSC threshold 0.143 | 9.9 | 10.9 | 4.5 | 17.9 | 12.2 | 11.4 | 26.4 |
| Map resolution range (Å) | 7.3-23.3 | 8-23.9 | 4.2-9.5 | 12.9-25.8 | 9.1-24.0 | 7.9-23.9 | 14.8-26 |

**Table 3:**
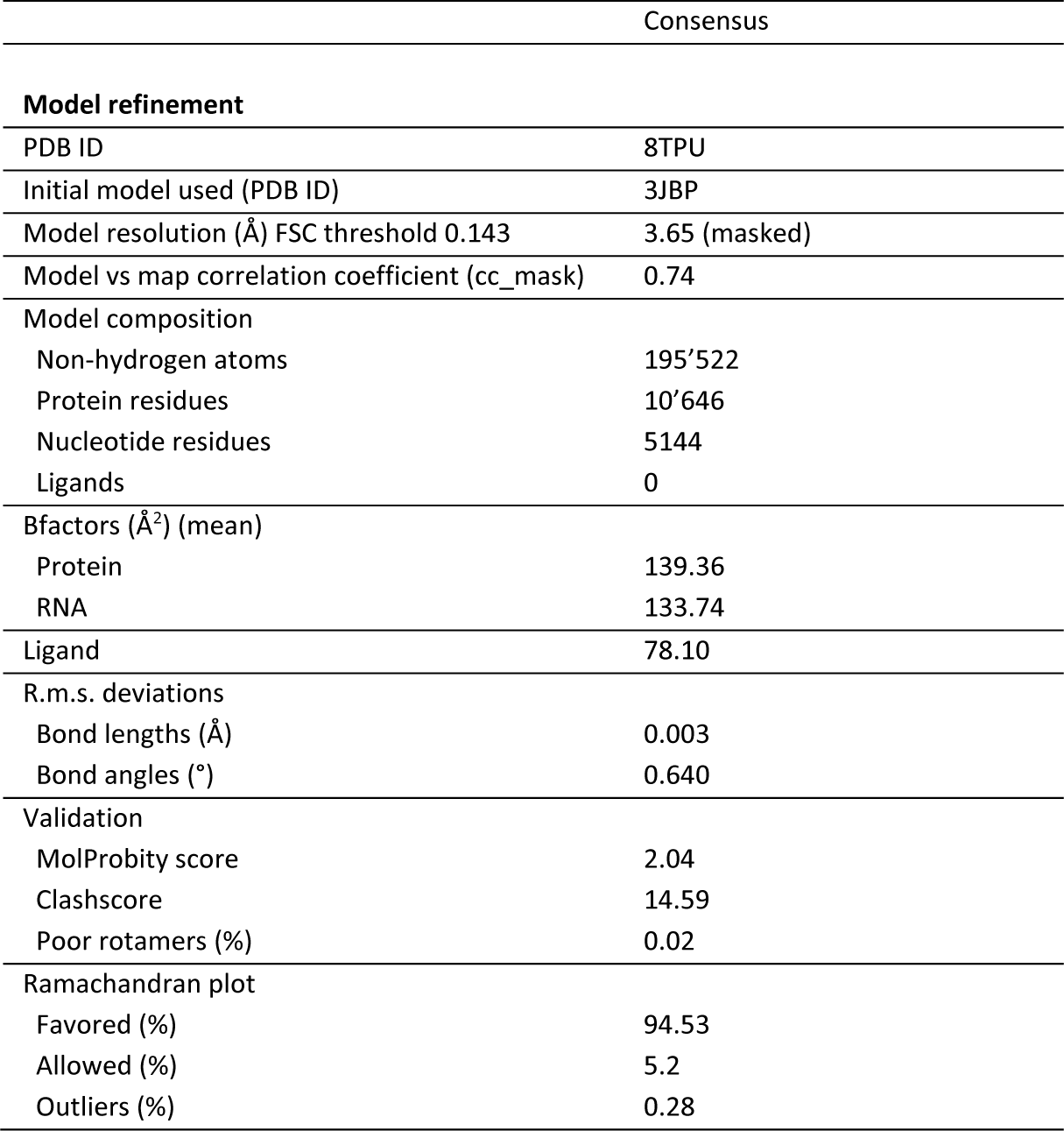
Model building and refinement.

**Extended Figure 1.**
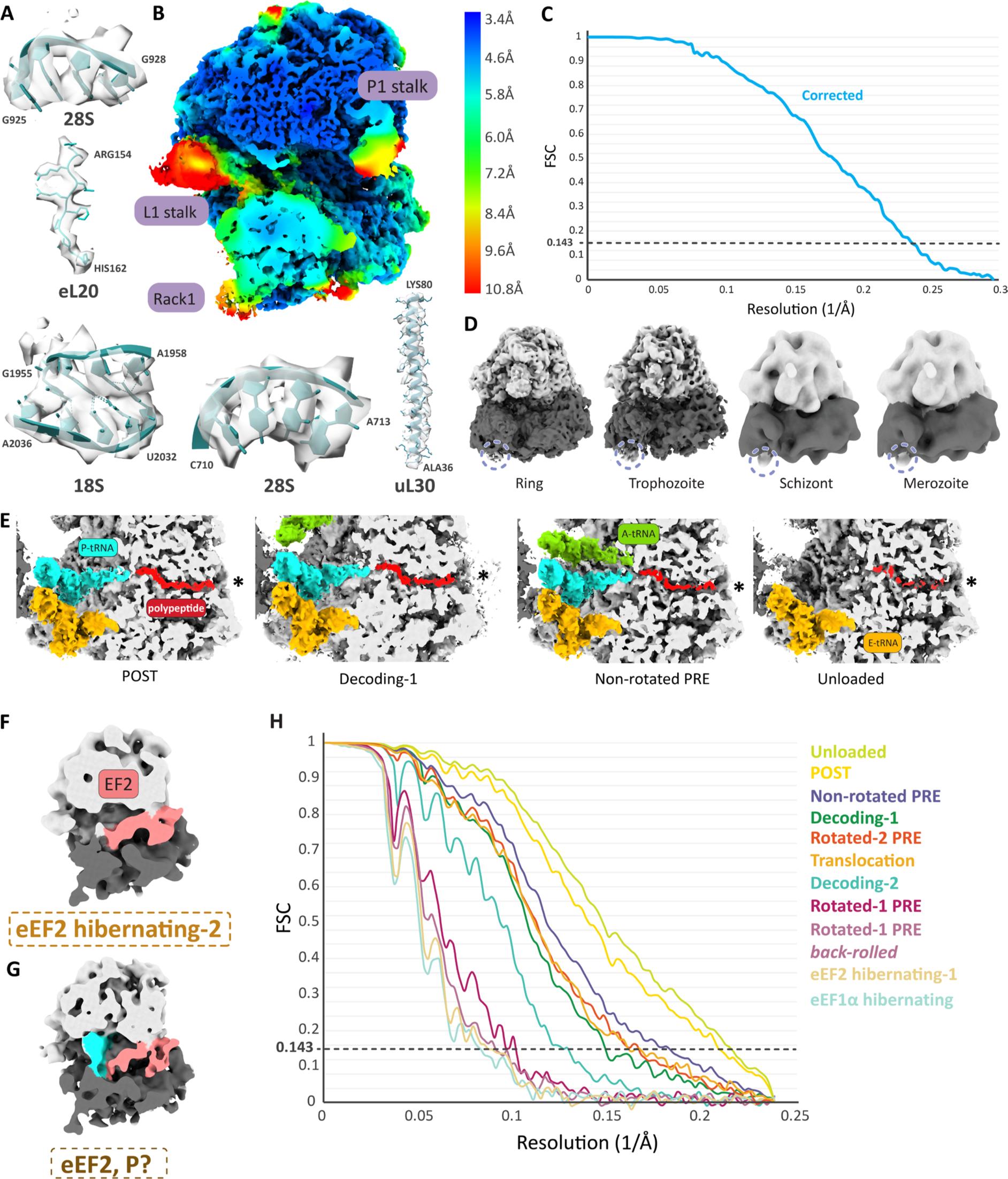
Detailed analysis of *Pf*80S ribosome features in consensus and resolution assessment of ribosomal states. **A**, Ribbon models of 28S and 18S rRNA and eL20 and uL30 ribosomal protein segments, shown with corresponding STA density (surface representation). **B**, Cross-section of *Pf*80S consensus map shown at lower threshold to improve clarity of low resolution features and colored according to local resolution, as calculated in RELION3.1. **C**, Fourier shell correlation (FSC) curve for the consensus reconstruction, calculated in RELION3.1 from comparison of two independently refined ‘half-maps’ from which global resolution is determined using the ‘Gold-standard’ 0.143 cut off (dotted line). **D**, *Pf*RACK1 density (lavender circle) is indicated in independent reconstructions of the *Pf*80S ribosomes from individual life cycle stages. **E**, Cross-section along the peptide exit tunnel of the POST, decoding-1, non-rotated PRE and unloaded *Pf*80S ribosomes. Density for tRNAs and nascent polypeptide chain (red) are shown as full volumes for each translation intermediate, and the distal end of the peptide exit tunnel is indicated with an asterisk. (60S=light grey, 40S=dark grey, A-site=green, P-site=cyan, E-site=orange). **F-G**, Cross-sections of eEF2 hibernating-2 (**F**) and unassigned eEF2, P? states (**G**). **H**, Fourier shell correlation (FSC) curves for ribosomal states identified through classification, calculated in RELION3.1 as described in (**C**). Identified ribosomal states are shown in descending order from highest to lowest resolution in the left side of the graph.

**Extended Figure 2.**
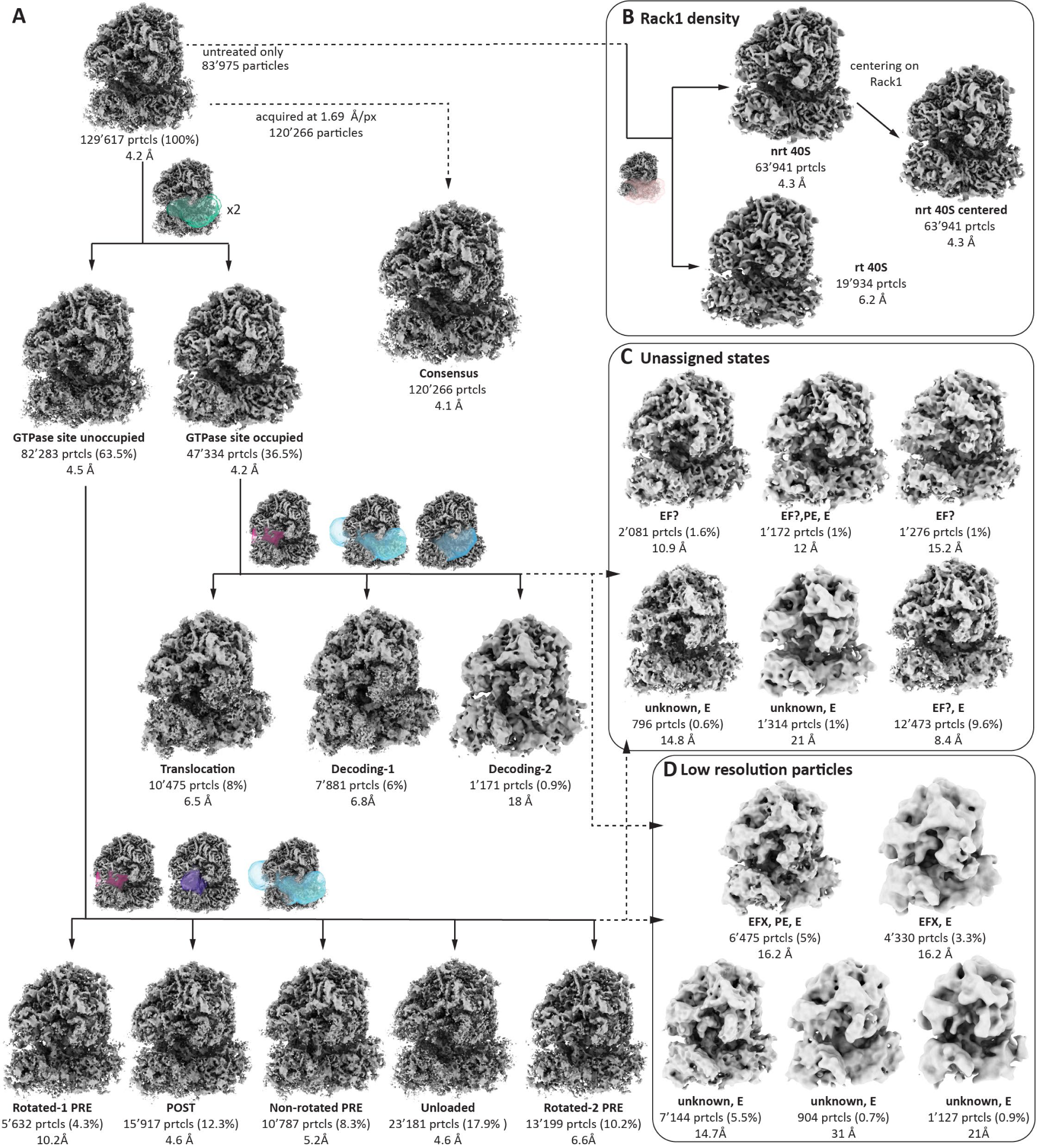
Focused classification of the subtomogram averaged *Pf*80S consensus reconstruction. **A,** Translation intermediates were separated following the displayed scheme using the following masks: elongation factor (EF) (green), PE-tRNA (magenta), EF-A-P-PE-E binding sites (cyan), A-tRNA (blue), P-tRNA (purple) (Fig. 3A, S3A-J). **B,** Processing strategy to improve *Pf*RACK1 density (Fig. 2D-M). **C,** Unassigned translational states with and without density in the EF site. **D,** Low resolution particles sorted out during focused classification. For all panels the number of particles, percentage of total particles and resolution are indicated below each map.

**Extended Figure 3.**
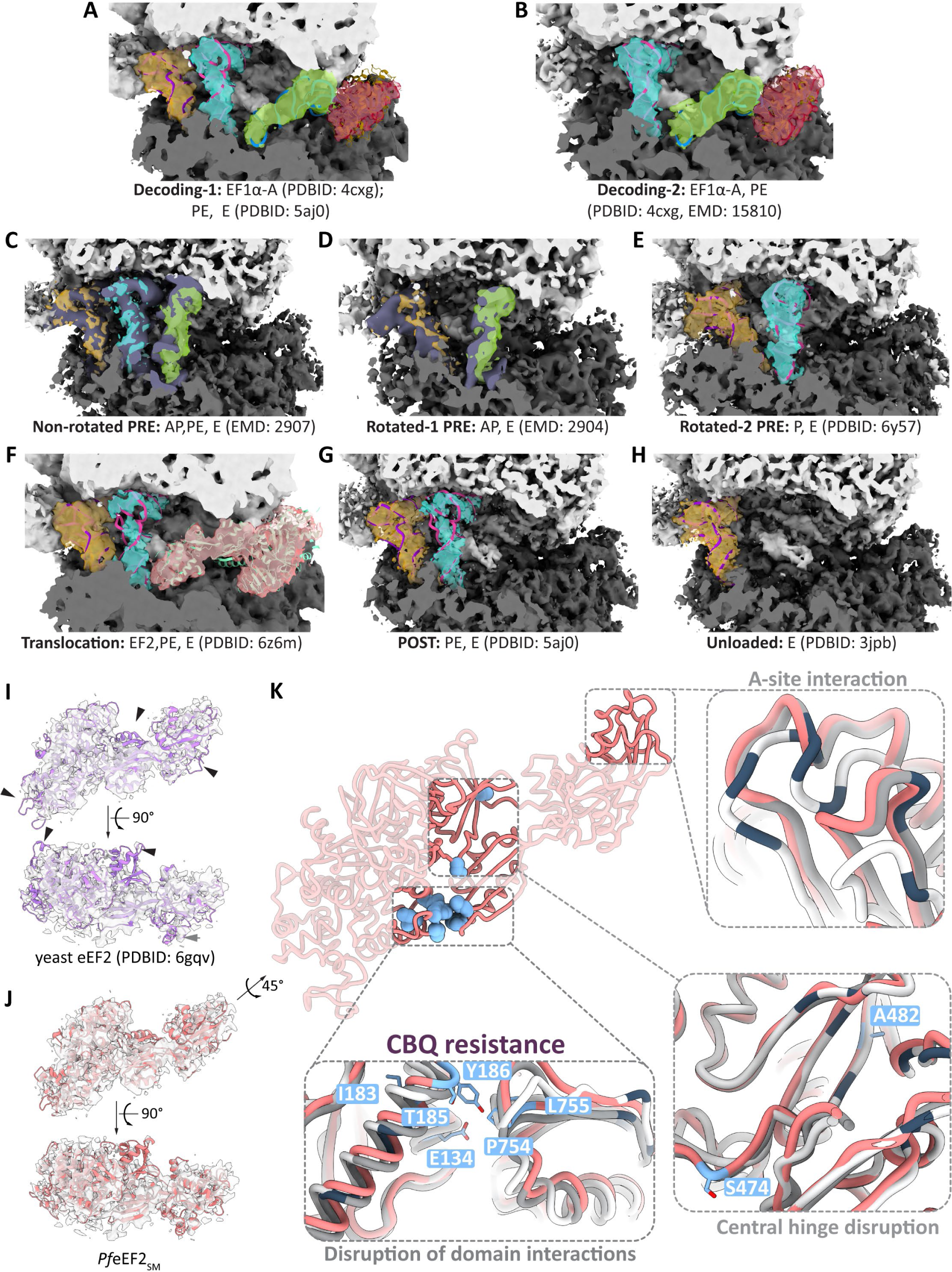
Identification of translation intermediates and analysis of elongation factor densities and models. **A-H**, Detailed view of the GTPase and tRNA binding sites in translation intermediate reconstructions, shown as cross-sections overlaid with published ribbon models or maps for assignment of states. Density of bound ligands is shown at 50% transparency for clarity. **I-J**, Ribbon model of yeast (purple) eEF2 solved in 80S context (**I**) and *Pf*eEF2_SM_ (SWISS-MODEL) (salmon) (**J**) fit into eEF2 density from our *Pf*80S translocation intermediate shown as side and 90°-rotated top views. **K**, Overview of mutations (light blue) conferring cabamiquine (CBQ) resistance in *P. falciparum* mapped onto *Pf*eEF2_SM_ (salmon) shown 45°-rotated from side view in (**J**). Insets show close up of affected regions overlaid with human (grey) and yeast (white) models. Reported human and yeast eEF2 mutations are shown in dark blue.

**Extended Figure 4.**
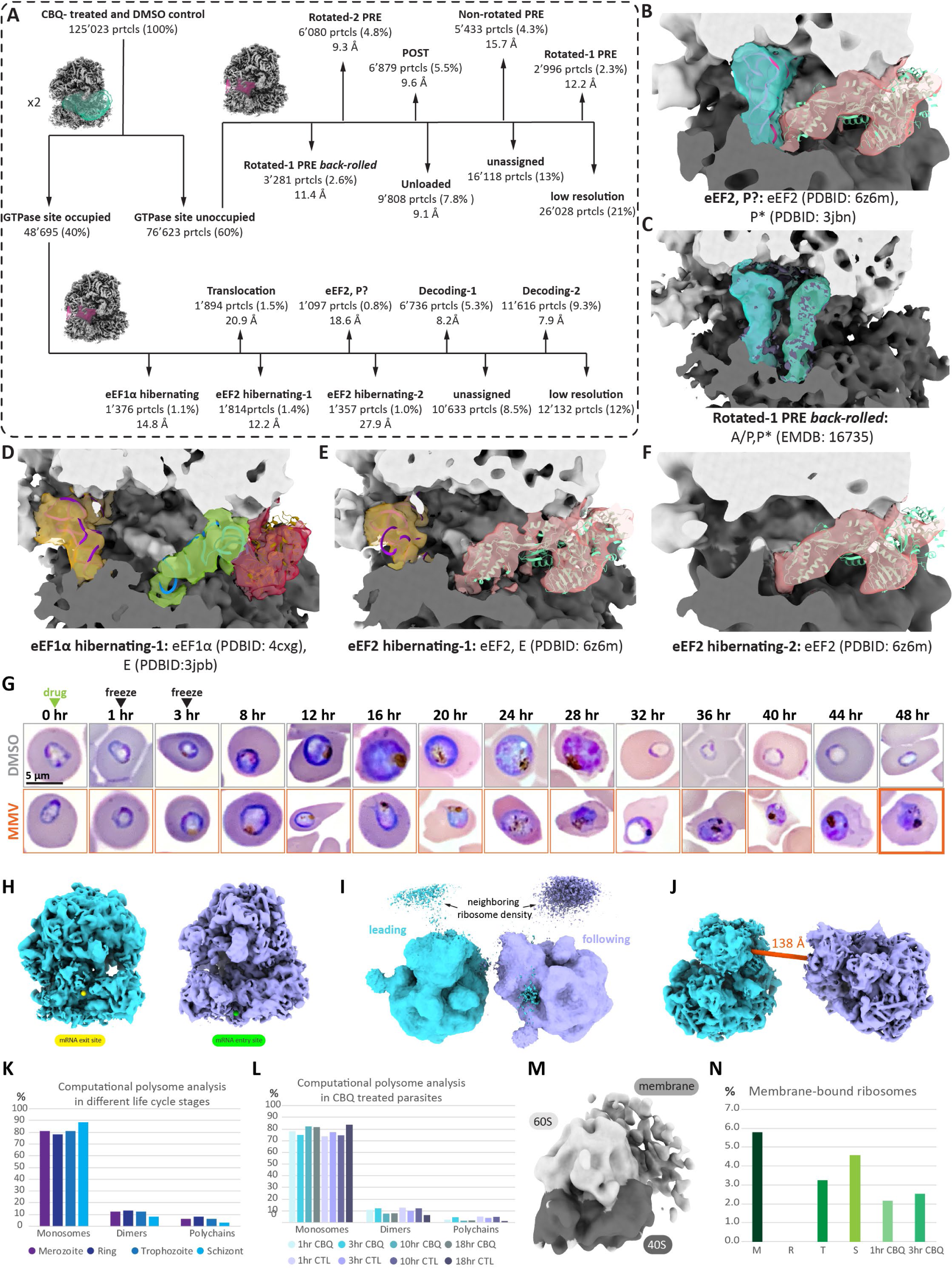
*Pf*80S ribosome analysis of CBQ-treated and DMSO-control parasites and 48 hour MMV019189-treatment time course. **A,** Classification scheme for translational states for CBQ-treated and DMSO control (CTL) parasites using masks for elongation factor (EF) (green) and PE-tRNA (magenta). Number of particles, percentage of total particles and resolution are indicated below name of translation intermediate. **B-H**, Detailed view of the GTPase and tRNA binding sites in reconstructions of additional translation intermediates, shown as cross-sections overlaid with published ribbon models or maps for assignment of states. Density of bound ligands is shown at 50% transparency for clarity. **G**, Giemsa-stained blood smears of highly synchronized early trophozoite-stage *P. falciparum* parasites, taken at 0, 1, and 3 hours after addition of DMSO or MMV019189 (MMV), and every 4^th^ hour after, out to 48 hours. Three biological replicates were performed, with **n= 6-80** for each time point in each replicate. One representative replicate is shown. Time of drug addition, as well as time points corresponding to those used in our *in situ* cryoET studies, are indicated with green and black arrowheads, respectively. MMV019189-treated parasites at 48 hr (bolded box) are 16.7-50% pyknotic. **H**, Position of mRNA exit (yellow sphere) and entry (green sphere) sites defined based on *Pf*80S ribosome reconstruction. **I**, Relative arrangement of two *Pf*80S ribosomes shown at lower threshold based on neighboring ribosome density, simulating leading and following ribosome in a polysome chain. **J**, Distance measurement in Å (ChimeraX, dark orange) between exit and entry site from leading to following ribosome. **K**, Percentage of monosomes, dimers and polychains across life cycle stages. **L**, Change in monosomes, dimers and polychains from CBQ treated-to DMSO-treated parasites across four time points. **M**, Reconstruction of *Pf*80S membrane-bound ribosomes (2’527 particles). **N**, Percentages of membrane bound *Pf*80S ribosomes across life cycle stages and CBQ treated parasites (M= Merozoite, R= Ring, T= Trophozoite, S= Schizont, CBQ= cabamiquine).

**Extended Figure 5.**
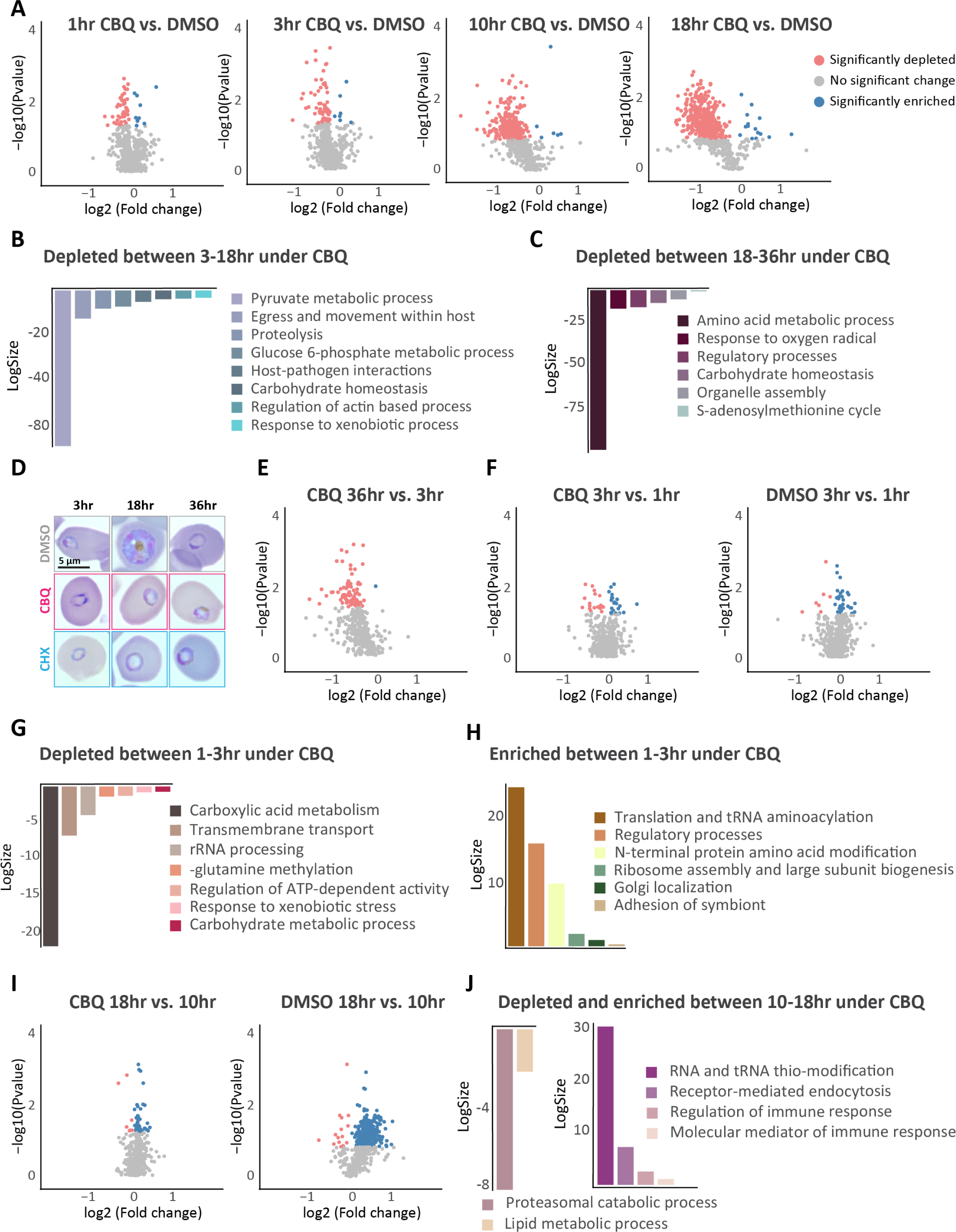
Quantitative proteomics analysis of consequences of translation inhibition in *P. falciparum*. **A,** Volcano plots showing differentially expressed proteins between CBQ- and DMSO-treated parasites at 1, 3, 10, and 18 hours post-treatment. B-C, Revigo-generated gene ontology (GO) enrichment analysis of depleted proteins under CBQ-pressure between 3-18 hours (B) and 18-36 hours (C). **D,** Giemsa-stained smears of DMSO-, CBQ- and CHX-treated parasites at 3, 18 and 36 hours. **E,** Volcano plot showing differentially expressed proteins under CBQ-pressure comparing proteins present at 36 hours to 3 hours. F,I, Volcano plots showing differentially expressed proteins under CBQ-pressure (left) and DMSO-control (right) comparing proteins present at 3 hours to 1 hour (F) and 18 hours compared to 10 hours (I). G-H, Revigo-generated GO enrichment analysis of specifically depleted (G) and enriched (H) proteins in CBQ-treated parasites at 3 hours compared to 1 hour. **J,** Revigo-generated GO enrichment analysis of specifically depleted (left) and enriched (right) proteins in CBQ-treated parasites at 18 hours compared to 10 hours. For all volcano plots, the mean average of each protein from three replicates (n=3) is shown. Y-axes show log10 p-value and x-axes show log_2_ fold change using nested study design. Dark coral and blue circles indicate proteins with significant enrichment and depletion, respectively. Grey circles indicate proteins with no significant (p>0.05) change in expression. For GO enrichment analysis graphs, bars are colored by Revigo-designated cluster. Y-axes are log_10_ of number of annotated GO term in *P. falciparum*. Each bar in the plot represents a sub-cluster.

**Extended Figure 6.**
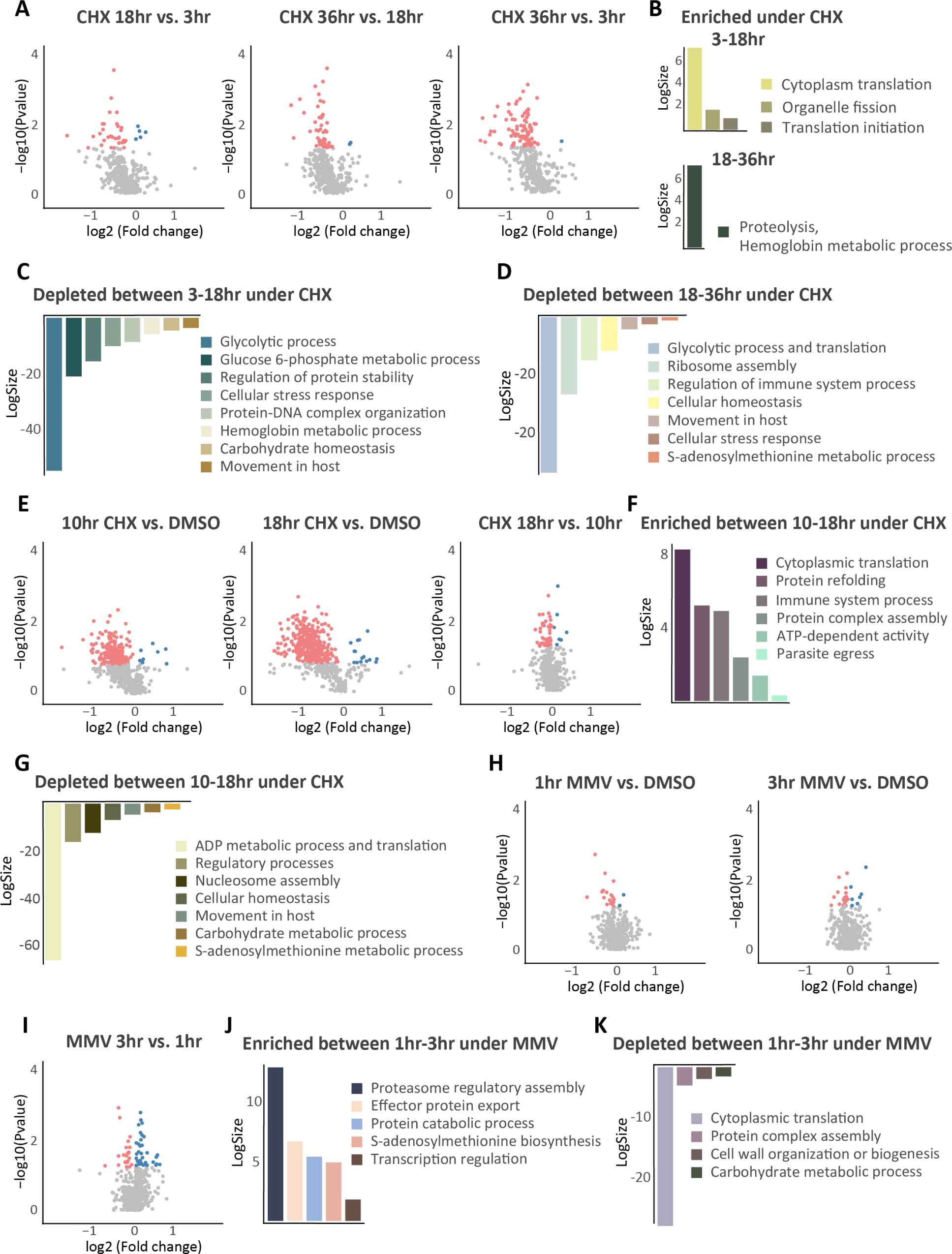
Quantitative proteomics analysis of consequences of translation inhibition in *P. falciparum*. **A**, Volcano plots showing differentially expressed proteins between different timepoints in CHX-treated parasites comparing 18 hours to 3 hours (left), 36 hours to 18 hours (middle) and 36 hours to 3 hours (right). **B-D**, Revigo-generated gene ontology (GO) enrichment analysis of enriched (B) and depleted **(C-D)** proteins in CHX-treated parasites at 18 hours compared to 3 hours (**B**, top, **C**) and 36 hours compared to 18 hours (**B**, bottom, **D**). **E,** Volcano plots showing differentially expressed proteins between CHX- and DMSO-treated parasites at 10 and 18 hours (left, middle) and comparing proteins present at 18 hours to 10 hours (right). **F-G**, Revigo-generated GO enrichment analysis of specifically enriched **(F)** and depleted **(G)** proteins in CHX-treated parasites at 18 hours compared to 10 hours. **H,** Volcano plots showing differentially expressed proteins between MMV- and DMSO-treated parasites at 1 (left) and 3 (right) hours. **I-K**, Volcano plots of differentially expressed proteins **(I)** and corresponding GO analysis of enriched **(J)** and depleted **(K)** under MMV-pressure between 1 and 3 hours. For all volcano plots, the mean average of each protein from three replicates (n=3) is shown. Y-axes show log10 p-value and x-axes show log_2_ fold change using nested study design. Dark coral and blue circles indicate proteins with significant enrichment and depletion, respectively. Grey circles indicate proteins with no significant (p>0.05) change in expression. For GO enrichment analysis graphs, bars are colored by Revigo-designated cluster. Y-axes are log_10_ of number of annotated GO term in *P. falciparum*. Each bar in the plot represents a sub-cluster.

**Extended Figure 7.**
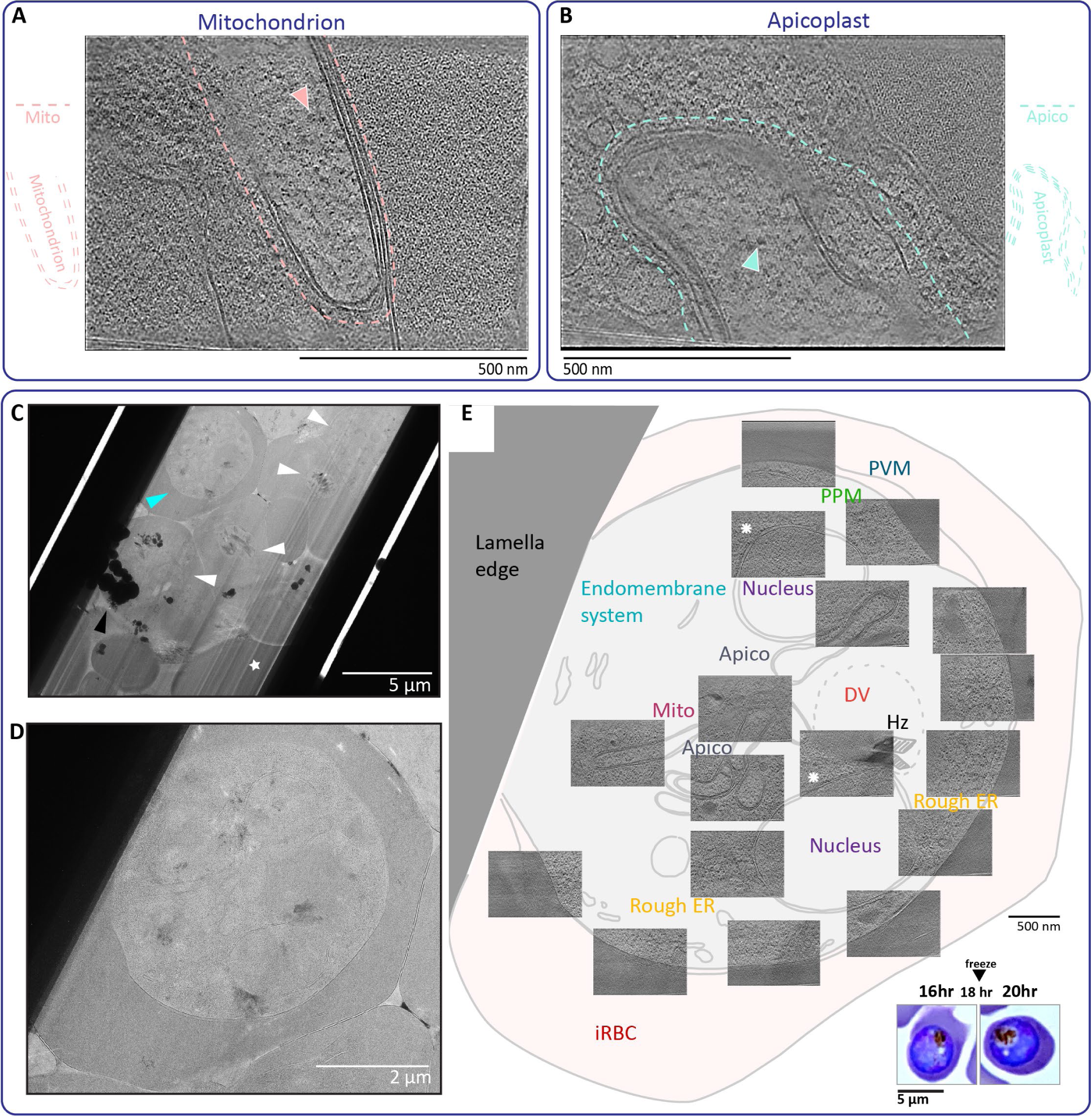
Details of parasite mitochondrion and apicoplast and 18 hr DMSO-control parasites. **A**, Representative tomogram illustrating parasite mitochondrion and mitochondrial ribosomes, indicated in pink trace and pink arrowhead, respectively. **B**, Representative tomogram illustrating parasite apicoplast and apicoplast ribosomes, indicated in cyan trace and cyan arrowhead, respectively. **C**, Montaged cryoTEM images of lamella containing DMSO-control parasites frozen, milled and imaged at 18 hours post-treatment. Parasites, contamination and milling artefacts are indicated with white arrow heads, black arrow heads and white stars, respectively. **D**, Close-up view of parasite indicated with cyan arrowhead, in the upper left of lamella in (**C**). **E**, Integrated cell montage view of cell in (**C**) created by overlaying averaged central slices of tomograms collected on cell in (**C**) on a tracing of the cell in (**C**). White asterisks indicate nuclear pore complexes. Giemsa-stain blood smears of DMSO-treated parasites smeared at 16 and 18 hours post-treatment are shown in the inset on the bottom right. PVM, parasitophorous vacuolar membrane. PPM, parasite plasma membrane. EMS, endomembrane system. DV, digestive vacuole. Hz, hemozoin. ER, endoplasmic reticulum. Mito, mitochondrion. Apico, apicoplast. iRBC, infected red blood cell.

